# Mitochondrial ROS-induced metabolic alterations differentially regulate ferroptosis sensitivity

**DOI:** 10.64898/2025.12.21.695848

**Authors:** Somesh Banerjee, Matthew Ryan Smith, Yining Irene Wang, Michael Wang, Derrik Gratz, Gregory K. Tharp, Sumin Kang, Peijian He

## Abstract

Ferroptosis is an iron-catalyzed lipid peroxidation (LP)-dependent cell death. Induction of mitochondrial ROS (mtROS) is crucial in the execution of ferroptosis, but the underlying mechanism remains unclear. Through utilizing the hepatocyte model and RNA-seq analysis, we determined mtROS-dependent metabolic changes that modulate ferroptosis sensitivity. Elevated mtROS production and LP suppressed glycolysis, fatty acid oxidation, and citric acid cycle activity, representing adaptive responses that protect cells from ferroptosis. On the other hand, mtROS-driven signaling impaired glutathione biosynthesis and downregulated genes involved in coenzyme Q10 (CoQ) biosynthesis, including those in the mevalonate pathway and CoQ8A, a key stabilizer of the CoQ biosynthetic complex. Importantly, silencing CoQ8A expression enhanced, whereas overexpression of CoQ8A reduced, ferroptosis susceptibility of hepatocytes and various cancer cell types. The mtROS-mediated downregulation of CoQ8A was dependent on farnesoid X receptor (FXR) and retinoid X receptors (RXRs). Collectively, our findings highlight that mtROS promotes ferroptosis, at least in part, by suppressing glutathione and CoQ biosynthesis.

## 1. Introduction

Hepatocytes play a pivotal role in metabolic processing and storage of the macronutrients including carbohydrates, fats, and proteins, as well as the micronutrients including iron, copper, and vitamins ^1–3^. Mitochondria are highly enriched in hepatocytes and are the central hub of cellular metabolism through the tricarboxylic acid (TCA) cycle, and oxidative phosphorylation that produces ATP. Mitochondria are also a primary site of reactive oxygen species (ROS) production. Mitochondrial dysfunction and oxidative stress constitute crucial mechanisms in liver disease progression in many contexts ^4^, including metabolic dysfunction-associated steatotic liver disease ^5^, alcohol-associated liver disease ^6^, viral hepatitis ^7^, and drug-induced liver injury ^8^. In the physiological state, mitochondrial ROS (mtROS) is produced at a low level in the electron transport chain and acts as signaling molecules influencing various cellular functions ^9^. However, overwhelming mtROS may be produced in pathological conditions due to elevations in nicotinamide adenine dinucleotide (NADH) and flavin adenine dinucleotide (FADH_2_) levels, mitochondrial iron overload, imbalances in oxidized/reduced coenzyme Q10 (CoQ) ratio, and mitochondrial membrane hyperpolarization ^10,11^. Excessive mtROS causes mitochondrial dysfunction, perturbations of cellular metabolism, and oxidative damages of the DNA, lipids and proteins that ultimately leads to oxidative cell death ^12^.

Ferroptosis is an iron-catalyzed lipid peroxidation (LP)-dependent necrosis resulting from an excess of labile iron in the context of compromised antioxidant defense against lipid ROS ^13^. The well-defined anti-LP mechanisms include the reduced glutathione (GSH)-GPx4 pathway and reduced form of CoQ (CoQH_2_, also known as ubiquinol), the production of which is mediated by ferroptosis suppressor protein (FSP1) ^14–16^, the activities of mitochondrial complex I/II ^17^, and dihydroorotate dehydrogenase (DHODH) ^18^. Thus, under conditions of reduced GPx4 activity and limited GSH or CoQH₂ availability, cells are predisposed to iron-driven, unrestrained LP, resulting in ferroptosis. Several pharmacological inducers of ferroptosis have been developed, including erastin, RSL3, iFSP1, and FIN56. Erastin reduces glutathione biosynthesis by inhibiting cystine transporter xCT ^13,19^, while RSL3 was believed to inactivate GPx4, despite recent evidence suggesting that RSL3 does not inhibit GPx4 activity ^20,21^. iFSP1 inhibits the activity of FSP1 restricting the availability of CoQH_2_ ^22^, and FIN56 induces both GPx4 protein degradation and CoQ depletion ^23^. In addition, acute iron overload also induces ferroptosis in multiple cell types, including hepatocytes and cardiomyocytes ^24–26^, both of which are highly enriched in mitochondria.

Compelling evidence, including our work, has suggested a crucial role of mtROS in mediating ferroptosis ^25,27,28^. Scavenging mtROS with mitochondria-targeted antioxidants abolishes ferroptosis by most, if not all, of the above listed inducers. Nevertheless, the precise mechanisms by which mtROS initiates or amplifies ferroptotic signal that ultimately leads to ferroptosis remain unclear. Previous studies have uncovered a strong interplay between metabolic activity and ferroptosis susceptibility ^29–32^. In this study, we utilized cultured hepatocytes and iron-induced mtROS production and ferroptosis as models to interrogate the effects of mtROS on hepatocellular metabolism as well as the impact of metabolic changes on hepatocyte susceptibility to ferroptosis. We found that iron induces mtROS-dependent suppression of glycolysis, fatty acid oxidation, TCA reactions, and mitochondrial respiration, serving as metabolic adaptations that protect against ferroptosis. Importantly, we also uncovered a novel mechanism whereby mtROS-dependent disruption of the mevalonate pathway and CoQ biosynthetic machinery reduces CoQ production, thereby increasing the sensitivity of hepatocytes to ferroptosis.

## 2. Materials and Methods

### 2.1. Reagents

The chemicals and reagents used in the current study are listed in Supplementary Table 1.

### 2.2. Animals

C57BL/6 mice purchased from the Jackson Laboratory were bred and housed at Emory University animal facility with temperature and humidity control. All mice were maintained under 12-h light and 12-h dark (light between 07:00 and 19:00) environment with water and food *ad libitum*. Male mice at 10-12 weeks of age were injected (i.p.) with 500 mg/kg acetaminophen (APAP) or saline as the vehicle control. After 6 h, mice were euthanized and livers were harvested. All animal procedures were approved by the Institutional Animal Care and Use Committee of Emory University.

### 2.3. Cell culture

Primary mouse hepatocytes (PMH) were isolated from male mice and cultured as previously described ^25^. Male PMH were used in this study for its greater responsiveness to ferroptosis induction, as demonstrated in our previous report ^24^. Briefly, mouse liver was pre-perfused with Leffert’s buffer (HEPES 10 mM, KCl 3 mM, NaCl 130 mM, NaH_2_PO_4_ 1 mM, glucose 10 mM, pH 7.4) containing EGTA, followed by perfusion with Liberase that consists of collagenase I and II. Dissociated hepatocytes were purified on a Percoll gradient, and cell viability is determined by staining with 0.02% trypan blue. PMH were seeded in collagen-coated plates for 3 h in William’s E medium supplemented with 5% FBS, 1x penicillin/streptomycin (P/S), 10 mM HEPES, 1x insulin and 40 ng/ml dexamethasone, followed by overnight culture in fresh medium containing 5% FBS, 1x P/S and 10 mM HEPES prior to any treatments. HepG2 cells, a human hepatoblastoma cell line, and H1299 cells, a human non-small-cell lung carcinoma cell line, were cultured in RPMI medium supplemented with 10% FBS and 1x P/S. MDA-MB-231 cells, a human breast cancer cell line, and MIA Paca-2 cells, a human pancreatic cancer cell line, were cultured in high-glucose DMEM supplemented with 10% FBS and 1x P/S.

### 2.4. Lentivirus production

Lentiviral expression constructs used in this study are listed in Supplementary Table 1. DNA constructs expressing shRNA targeting Coq8a, Nr1h4 (Fxr), Hmgcs1, and Rxra were purchased from Sigma-Aldrich. Constructs that express V5-tagged (C-terminus) COQ8A, CPT1A, and PDHA1 were obtained from DNASU plasmid repository. Lentiviral particles, prepared as previously described ^24^, were utilized for cell infections in the presence of 5 μg/ml polybrene. Stable expression was achieved by selecting the infected cells with 5 μg/ml puromycin (for knockdown) or 5 μg/ml blasticidin (for overexpression). Selection antibiotics were excluded whenever cells were seeded for experiments.

### 2.5. Live cell imaging

Propidium iodide (PI) staining of the cultured cells was performed as previously described ^24,25^. Briefly, treated and untreated hepatocytes were incubated for 30 min with 2 μg/ml PI in the culture medium, followed by fluorescent imaging with the Olympus IX83 imaging system. The numbers of PI^+^ cells and total numbers of cells *per* field were counted, from which the percentage of PI^+^ cell death was calculated. Intracellular neutral lipids were stained with 2 μM BODIPY 493/503. LP was assessed by incubating cultured cells with 2 μM BODIPY 581/591 C11. After washing with PBS, cells were imaged at the emissions of ∼590 nm (red) and ∼510 nm (green). The intensity of green fluorescence signal indicates the extent of LP.

### 2.6. Quantitative reverse transcription PCR (qRT-PCR) analysis

Total RNA was extracted from cultured cells using the RNeasy Mini Kit. Two μg of total RNA was used for first strand cDNA synthesis using the High-Capacity cDNA Reverse Transcription Kit according to the manufacturer’s instruction. Quantitative PCR was performed with *Power* SRBR Green PCR Master Mix on Bio-rad CFX Opus PCR system. PCR primer sequences are listed in Supplementary Table 2.

### 2.7. RNA sequencing (RNA-seq) analysis

PMH were pretreated for 30 min with or without 5 μM liproxstatin-1 (Lip-1), a lipid ROS scavenger and ferroptosis inhibitor, or 10 μM Mito-TEMPO (MT), a mitochondria-targeted antioxidant. Following pretreatment, both the pretreated and one group of untreated PMH were exposed to 100 μM ferric ammonium citrate (FAC) for 4 h. Cells were then lysed in 350 μL of Buffer RLT Plus and extracted using the RNeasy Micro kit (Qiagen) with on-column DNase digestion. RNA quality was assessed using a TapeStation 4200 (Agilent) and then ten nanograms of total RNA was used as input for cDNA synthesis using the Clontech SMART-Seq v4 Ultra Low Input RNA kit (Takara Bio) according to the manufacturer’s instructions. Amplified cDNA was fragmented and appended with dual-indexed barcodes using the Nextera XT DNA Library Preparation kit (Illumina). Libraries were validated by capillary electrophoresis on a Fragment Analyzer (Agilent), pooled at equimolar concentrations, and sequenced with PE100 reads on an Illumina NovaSeq 6000, yielding ∼35 million reads per sample on average. Alignment was performed using STAR version 2.7.9a and transcripts were annotated using the GRCm38 mouse genomic assembly and annotation. Transcript abundance estimates were calculated internal to the STAR aligner using the algorithm of htseq-count. DESeq2 was used for normalization and differential expression analysis using the Wald test. Functional enrichment was performed using the Gene Set Enrichment Analysis (GSEA) method implemented in the fgsea R package against genesets in collections from the Molecular Signatures Database (MSigDB). Log2 fold changes are calculated relative to the median expression in the control group. The p-values for significantly differentially expressed genes and pathways are multiple-test corrected via the Benjamini-Hochberg method. The data has been deposited with the NCBI GEO with the accession number GSE302789.

### 2.8. Western blotting

Cultured cells were rinsed with cold PBS and then lysed in 1x lysis buffer containing 20 mM Tris-HCl (pH 7.5), 150 mM NaCl, 1 mM β-glycerophosphate, 2.5 mM sodium pyrophosphate, 1 mM Na_2_-EDTA, 1 mM EGTA, 1 mM Na_3_VO_4_, 1 μg/ml leupeptin, 1% Triton X-100, and 1x Halt protease/phosphatase inhibitors. Protein supernatants were obtained following sonication and centrifugation, and protein concentration was determined by the Bicinchoninic Acid (BCA) Protein Assay. Protein lysates were boiled at 95 °C for 10 min in 1x Laemmli buffer, separated on SDS-PAGE gel, and transferred onto nitrocellulose membrane for Western blotting with the antibodies listed in Supplementary Table 1. Densitometric analysis was performed by using the ImageJ software.

### 2.9 Glycolysis assay

The glycolysis rate was determined by measuring the L-lactate concentration using a Glycolysis Cell-Based Assay Kit. Briefly, PMH were seeded in 12-well plates at the density of 2 x 10^5^ cells *per* well. On the next day of culture, following a change with fresh medium (no phenol red), cells were treated with FAC, 2-Deoxy-D-Glucose (2-dG), or rotenone for different time periods. Culture medium was then collected for measuring lactate content with a plate reader following the manufacturer’s instruction.

### 2.10. Glutathione assay

Reduced GSH levels and GSH/GSSG ratio were measured using a Glutathione Assay Kit. Briefly, PMH seeded in 6-well plates at the density of 4 x 10^5^ cells *per* well. Following treatments with FAC or rotenone, cell lysates were collected and processed to determine total and oxidized glutathione (GSSG) levels. GSH was calculated using the formula: GSH = total glutathione – 2 x GSSG, and results were expressed as nmol/mg protein.

### 2.11. Analysis of Mitochondrial Bioenergetics

Mitochondrial function of hepatocytes was assessed using extracellular flux analysis (Agilent; Santa Clara, CA). PMH were seeded onto XF96 microplates at a cell density of 5 x 10^3^ cells *per* well. Cells were treated with or without FAC, Lip-1 or MT, and then incubated for 1 h in XF DMEM assay media (DMEM supplemented with 10 mM glucose, 2 mM glutamine, 1 mM pyruvate, and buffered with HEPES, pH 7.4). Extracellular flux analysis measures oxygen consumption rate (OCR) at baseline, followed by sequential injection of the inhibitors oligomycin (1 μM; 25μL), carbonyl cyanide-4-trifluoromethoxyphenylhydrazone (1 μM; 25μL), and antimycin A/rotenone (0.5 μM; 25μL). At the end of the assay, protein abundance was measured using the BCA protein assay. The Agilent Seahorse Wave Pro Desktop software was used for processing and analyzing the raw data, and the results were expressed as pmol O_2_ consumed/min/ng protein.

### 2.12. Statistical analyses

Data were analyzed with the GraphPad Prism software (GraphPad Software, La Jolla, CA, USA) and are presented as mean ± SEM. Statistical significance was determined by unpaired *t*-test, with a value of *P* < 0.05 was considered significant.

## 3. Results

### 3.1. mtROS promotes hepatocyte ferroptosis through time-course dependent effects

We and others have previously demonstrated that pretreatment with mitochondrial ROS scavengers can prevent ferroptosis in response to inducers ^25,27,28^. In this study, we have provided additional evidence that pretreating HepG2 cells with rotenone, an inducer of mtROS *via* inhibition of mitochondrial complex I, significantly enhanced ferroptosis by potent inducers including FAC and RSL3 (**Fig. 1A**). We further observed that the percentage of PI+ cells was markedly higher in HepG2 cells (**Fig. 1B, *top***) and PMH **Fig. 1B, *bottom***) treated with FAC for 24 h compared to 4 h. Our previous findings showed that FAC treatment resulted in mitochondrial iron loading and mtROS production within a few hours ^25^. We hypothesize that FAC-induced mtROS triggers yet unrecognized molecular changes in the early phase, promoting ferroptosis during the later phase of exposure to FAC. To test this hypothesis, we first treated HepG2 cells and PMH with FAC for 4 h, followed by co-treatment with mitochondria-targeted antioxidant MT, for an additional 20 h. Although this treatment regimen reduced ferroptosis compared with the 24-h FAC treatment alone, it resulted in a significantly higher percentage of PI+ cells than the 4-h FAC treatment group (**Fig. 1B**). In contrast, substituting MT with Lip-1, a lipid ROS scavenger and ferroptosis inhibitor, prevented the increase over the 4-h FAC treatment group (**Fig. 1B**). Notably, when administered prior to FAC treatment, MT and Lip-1 were comparably effective in reducing FAC-induced ferroptosis (**Fig. 1B**). These findings suggest that, once mtROS-associated signaling is initiated, scavenging mtROS is less effective at preventing ferroptosis than directly attenuating LP. In the following sections, we aimed at identifying mtROS-dependent cellular changes that impact hepatocyte sensitivity to ferroptosis.

**Figure 1:**
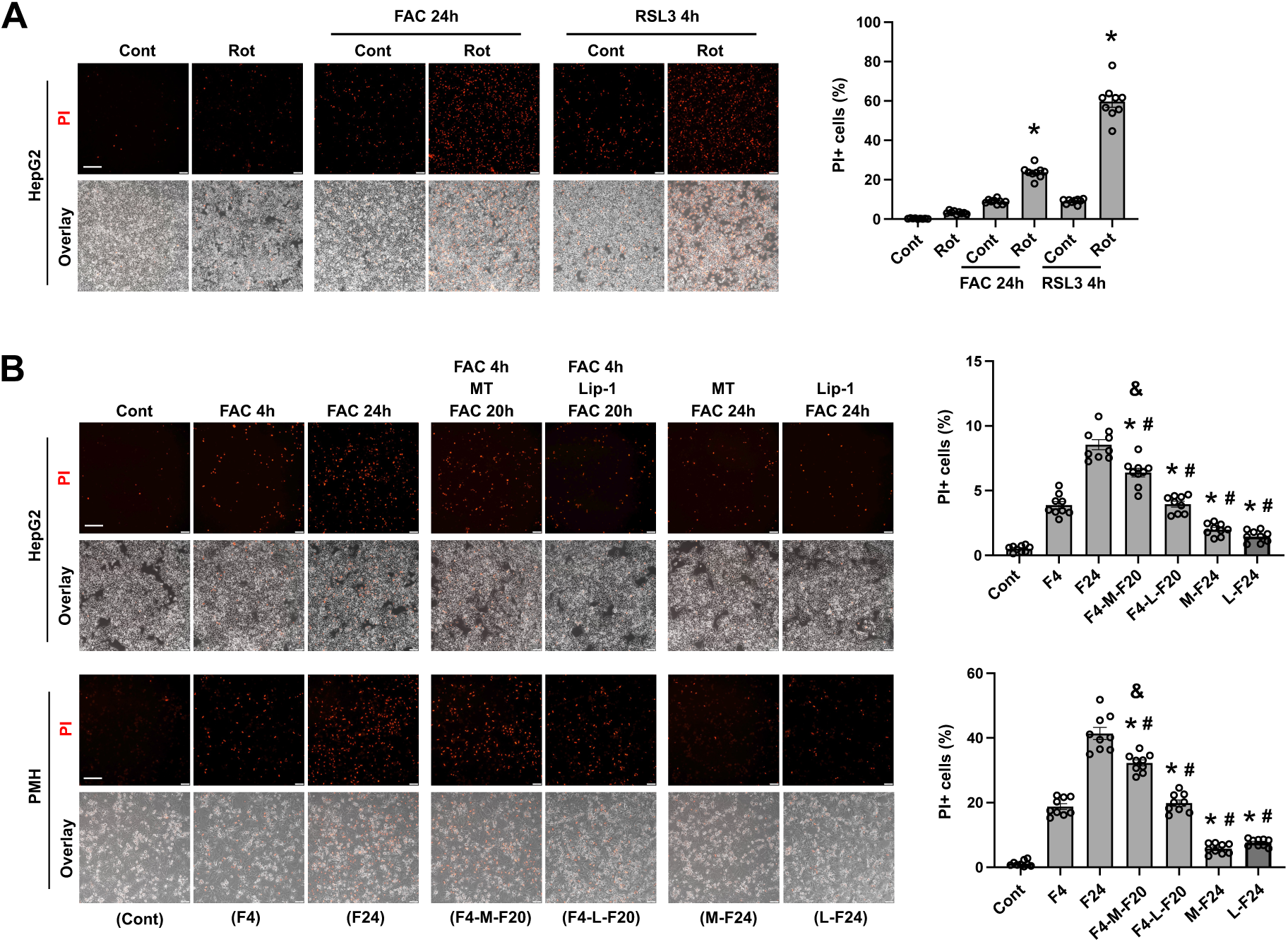
mtROS promotes ferroptosis of cultured hepatocytes. (**A**) HepG2 cells were pretreated for 4 h with or without 100 nM rotenone, followed by treatment with ferric ammonium citrate (FAC, 100 μM) for 24 h or 100 nM RSL3 for 4 h. Ferroptotic cells were stained with 2.5 μg/ml PI in the culture medium. (**B**) HepG2 cells and PMH were pretreated for 30 min with Mito-TEMPO (MT, 10 μM) or liproxstatin-1 (Lip-1, 5 μM) prior to treatment with FAC for 24 h. Alternatively, HepG2 cells and PMH were treated with MT or Lip-1 following 4-h FAC treatment and then stained with PI after additional 20 h. The percentage of PI+ cells are shown as mean±SEM (n = 9 fields of 3 independent experiments). *, *P* < 0.01 compared to untreated control (Cont) within the same group; ^#^, *P* < 0.01 compared to cells treated with 24-h FAC alone; and ^&^, *P* < 0.01 compared to cells treated with 4-h FAC alone. *Bars*, 200 μm.

### 3.2 Iron overload induces mtROS-dependent alterations in redox-active metal transport

To understand mtROS induced specific changes that regulate hepatocyte vulnerability to ferroptosis, we performed bulk RNA-seq analysis on PMH treated with FAC, with or without the addition of MT or Lip-1. Compared to untreated controls, FAC treatment led to the upregulation of 554 genes and downregulation of 458 genes with a log2 fold change threshold > 0.5 (**Fig. S1A**). Of note, only a small number of genes were differentially expressed between the “MT+FAC” and “Lip-1+FAC” groups (**Fig. S1B**), suggesting largely overlapping effects of mtROS and LP-mediated signaling on iron-induced transcriptional changes. This finding is consistent with previous reports, including our own, that mtROS scavengers effectively block LP-dependent ferroptosis ^25,27,28^.

Next, we analyzed changes in genes involved in hepatocellular iron metabolism. As expected, iron overload induced a significant reduction in the mRNA expression of transferrin-bound iron importer 1 (TfR1/TfRc) and an increase in the expression of iron exporter ferroportin (Slc40a1) (**Fig. 2A**). Divalent metal transporter 1 (Dmt1/Slc11a2), another iron importer expressed on both the plasma membrane and endosomal membrane ^33^, was only modestly decreased (**Fig. 2A**). The mRNA expression of heavy (Fth1) and light (Ftl) chain ferritins, iron binding proteins ^34^, was significantly increased in response to intracellular iron loading (**Fig. 2A**). Intriguingly, iron-induced elevation in ferritin transcript levels was attenuated by pretreatment with MT (**Fig. 2A**), consistent with previous findings of Nuclear Factor Erythroid 2-Related Factor 2 (NRF2), a key transcription factor in antioxidant defense, in redox-sensitive transcriptional regulation of ferritin ^35^. In contrast, iron-induced changes in TfR1 and ferroportin was not dependent on mtROS or LP.

**Figure 2:**
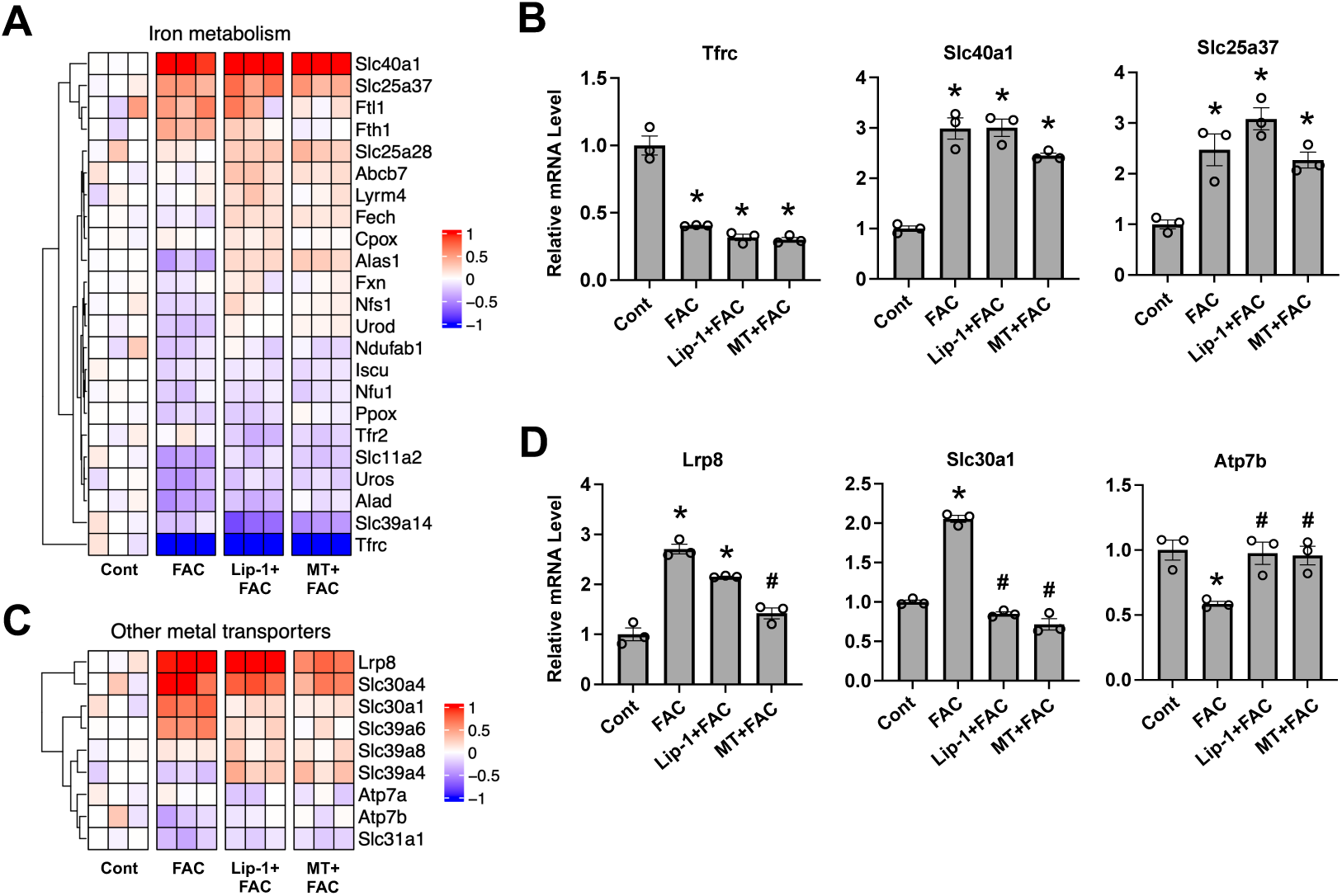
Iron induces mtROS and lipid peroxidation dependent changes in the expression of redox-active metal transporters. Cultured PMH were pretreated with or without 5 μM Lip-1 or 10 μM MT prior to treatment for 4 h with or without 100 μM FAC. Cells were harvested for bulk RNA-seq and qRT-PCR analyses. (**A**) Heatmap showing the log_2_ fold changes in mRNA expression of genes involved in iron metabolism pathway. (**B**) qRT-PCR analysis of mRNA expression levels of Tfrc, Slc40a1, and Slc25a37. (**C**) Heatmap showing the log_2_ fold changes in mRNA expression of genes responsible for copper, zinc and selenium transport. (**D**) qRT-PCR analysis of mRNA expression levels of Lrp8, Slc30a1, and Atp7b. Data of qPCR analysis are shown as mean±SEM (n = 3). *, *P* < 0.01 compared to untreated Cont; ^#^, *P* < 0.01 compared to cells treated with FAC alone.

Mitochondria are the primary site of iron utilization for the biosynthesis of iron-sulfur cluster (ISC) and heme – crucial cofactors for proteins involved in electron transport and cellular metabolism ^36,37^. We found that Mfrn1 (Slc25a37), a mitochondrial inner membrane iron importer ^38^, was upregulated by iron overload independently of mtROS or LP signaling (**Fig. 2A**). The mRNA expression of Fxn, Iscu and Nfs1, key genes involved in ISC biogenesis ^39^, was not significantly affected by iron overload (**Fig. 2A**). However, several genes in the heme biosynthesis pathway ^40^, including Alas1, Alad, and Uros, were downregulated in the context of iron overload (**Fig. 2A**). These changes in Tfr1, ferroportin, and Mfrn1 were further validated by qRT-PCR analysis (**Fig. 2B**).

Copper, a transitional metal, can also catalyze ROS production. We found that iron loading induced a slight reduction in the mRNA expression of Ctr1 (Slc31a1) and Atp7b (**Fig. 2C**), responsible for copper uptake and export in hepatocytes ^41^, respectively. In contrast to iron and copper, zinc is generally considered to play an antioxidant role by serving as a cofactor for many enzymes in antioxidant defense ^42^. Surprisingly, iron overload led to upregulation of zinc exporters Znt1 (Slc30a1) and Znt4 (Slc30a4), along with an increase in the expression of zinc importer Zip6 (Slc39a6) (**Fig. 2C**). Selenium, a redox-active trace element, exerts antioxidant functions through direct radical scavenging and as a cofactor in GPx enzymes ^43^. We observed a significant upregulation of Lrp8, a key selenium importer ^44^, in response to FAC treatment (**Fig. 2C**). These iron-induced changes in the expression of copper, zinc and selenium transporters were partially dependent on mtROS signaling and were confirmed by qRT-PCR (**Fig. 2D**).

### 3.3 Iron induces mtROS-dependent and -independent antioxidant responses

We previously showed that NRF2 protein expression was significantly upregulated in PMH following 4 h of FAC treatment, but GPx4 protein level was only increased at a later time point ^24^. Our RNA-seq analysis did not reveal significant changes in the mRNA expression of Nfe2l2 (encodes NRF2) or Gpx4 in PMH treated with FAC for 4h (**Fig. 3A**). The transcriptional activity of NRF2 depends on heterodimerization with small Maf proteins, including MafF, MafG, and MafK ^45^. Interestingly, the mRNA expression of MafF, MafG, and MafK was significantly increased in response to iron overload (**Fig. 3A**). Iron overload also significantly increased the mRNA expression of Atf4 and Atf3, transcription factors that are activated by, and confer protection against, various stress signals including oxidative stress ^46,47^. The expression of Hmox1, which encodes heme oxygenase-1 (HO-1), an antioxidant enzyme that mediates heme degradation ^48^, was robustly upregulated by iron overload. Notably, the induction of the above listed genes, except for Hmox1, was largely dependent on mtROS and LP-associated signaling, albeit to varying degrees (**Fig. 3A**). In contrast, we did not observe significant changes in the mRNA expression of other redox regulators, including thioredoxins (Txnrd2, Txn1, Txn2), superoxide dismutases (Sod1, Sod2), and hydrogen peroxide-scavenging enzymes such as peroxiredoxins (Prdx) and catalase (Cat) (**Fig. 3A**), within the examined duration of FAC treatment.

**Figure 3:**
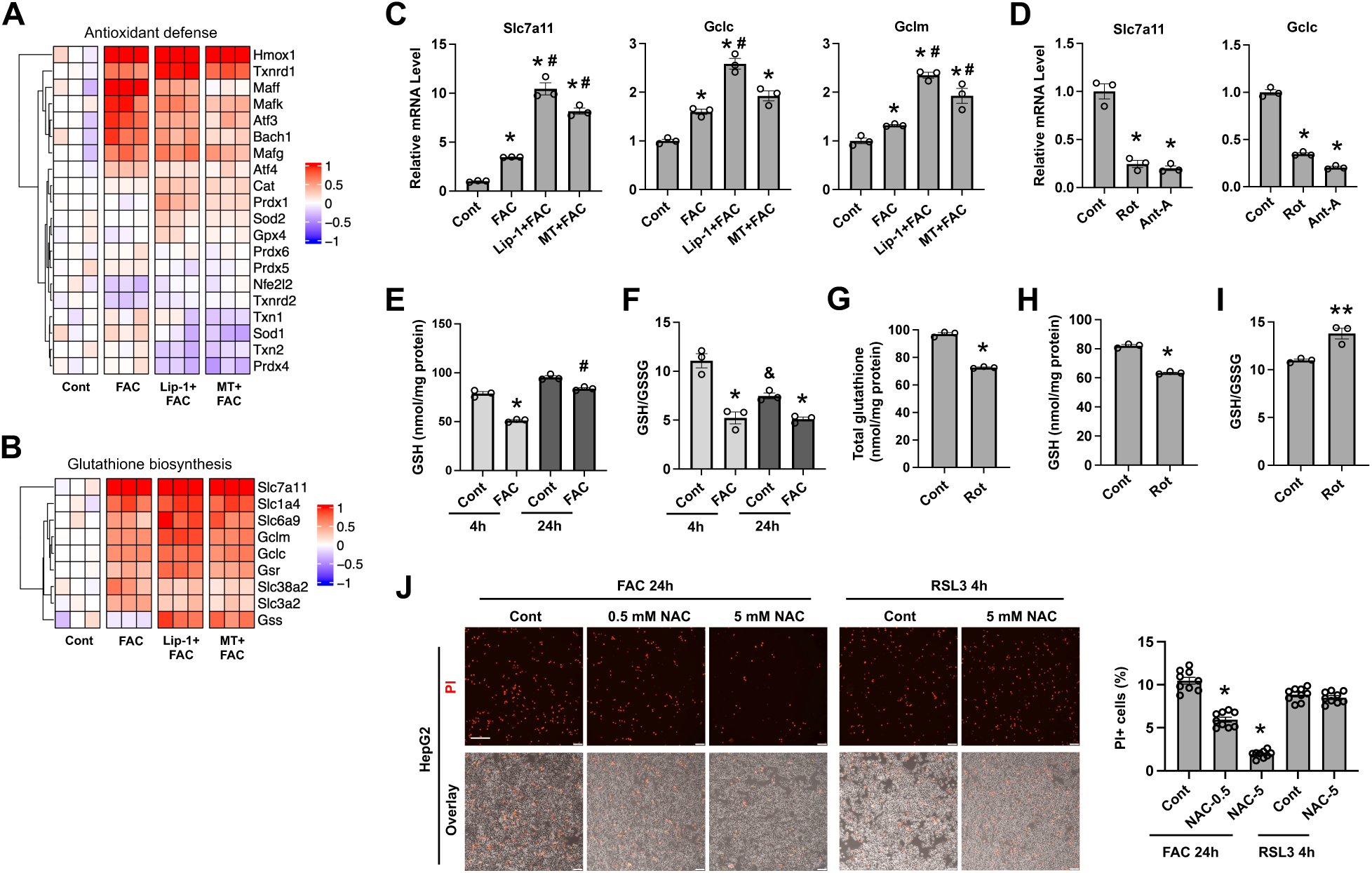
Iron overload induces mtROS-dependent and -independent oxidative stress responses. PMH were pretreated with or without 5 μM Lip-1 or 10 μM MT prior to treatment for 4 h with or not 100 μM FAC, followed by RNA-seq and qRT-PCR analyses (**A-C**). (**A**) Heatmap showing the log_2_ fold changes in mRNA expression of genes involved in antioxidant defense. (**B**) Heatmap showing changes in mRNA expression of genes involved in glutathione biosynthesis. (**C**) qRT-PCR analysis of mRNA expression levels of Slc7a11, Gclc, and Gclm. (**D**) Slc7a11 and Gclc mRNA expression in PMH treated for 4 h with or without 200 nM rotenone (Rot) or 10 μM antimycin A (Anti-A). Reduced form glutathione (GSH) content (**E**) and GSH/GSSG ratio (**F**) in PMH treated with or without 100 μM for 4 h or 24 h. Total glutathione (**G**) and GSH (**H**) contents and GSH/GSSG ratio (**I**) in PMH treated for 4 h with or not 200 nM Rot. (**J**) HepG2 cells were pretreated for 4 h with or without 0.5 mM or 5 mM N-acetylcysteine (NAC), followed by treatment for 24 h with 100 μM FAC or 4 h with 100 nM RSL3. Necrotic cells were stained with PI, and the percentage of PI+ cells are shown as mean±SEM (n = 9 fields of 3 independent experiments). *, *P* < 0.01 and **, *P* < 0.05 compared to Cont within the same group; ^#^, *P* < 0.01 compared to cells treated with FAC for 4 h; and ^&^, *P* < 0.01 compared to Cont at the 4-h time point. *Bar*, 200 μm.

Glutathione is a non-enzymatic antioxidant that plays a central role in redox homeostasis, particularly in GPx4-dependent defense against LP. Glutathione biosynthesis requires cysteine, glycine, and glutamate as substrates and is catalyzed by glutamate-cysteine ligase (GCL) and glutathione synthetase (GS) ^49^. Our data showed that the glutathione biosynthesis pathway was significantly activated in response to iron overload (**Fig. 3B**). Specifically, we observed increased mRNA expression of several amino acid transporters required for glutathione synthesis, including the cystine/cysteine transporters Slc7a11, Slc3a2, and Slc1a4, the glycine transporter SLC6A9, and the glutamine transporter SLC38A2 (**Fig. 3B**). The expression of Gclc and Gclm, which encode the catalytic and regulatory subunits of GCL, was also elevated, whereas Gss, encoding GS, was not significantly affected (**Fig. 3B**). Glutathione reductase (GSR) mediates the regeneration of reduced glutathione (GSH) from the oxidized form glutathione disulfide (GSSG). We showed that Gsr mRNA level was also increased by FAC treatment (**Fig. 3B**). Of note, except for Slc3a2 and Slc38a2, the expression of most genes in the glutathione pathway was upregulated by iron in an mtROS-independent manner (**Fig. 3B**). The iron-induced upregulation of Slc7a11, Gclc, and Gclm was confirmed by qRT-PCR (**Fig. 3C**). Interestingly, treatment with either MT or Lip-1 further enhanced the expression of these genes. Notably, a more pronounced increase in Slc7a11 and Gclc expression was observed following 24 h of FAC treatment (**Fig. S2**). In contrast to iron overload, which induces ROS production in mitochondria and extra-mitochondrial compartments, rotenone and antimycin A, potent inducers of mtROS, led to marked reductions in the expression of Slc7a11 and Gclc (**Fig. 3D**).

We hypothesized that iron catalyzed ROS production leads to glutathione depletion, thereby triggering upregulation of the glutathione biosynthetic pathway as a compensatory response. Indeed, both total glutathione levels (**Fig. 3E**) and the GSH/GSSG ratio (**Fig. 3F**) were significantly reduced in PMH after 4 h of FAC treatment. After 24 h, total glutathione content increased relative to the 4-h time point, whereas the GSH/GSSG ratio remained significantly lower than in untreated controls (**Fig. 3E-F**). An elevation in total glutathione levels with prolonged FAC treatment is likely ascribed to increased expression of amino acid transporters, including SLC7A11 and SLC3A2. Additionally, iron overload significantly decreased the expression of many genes of the cytochrome P450 family that participate in xenobiotic metabolism (**Fig. S3**). As these enzymes consumes NADPH and produces ROS, this change likely represents another adaptive response of the already-stressed hepatocytes. In line with the downregulated expression of Slc7a11 and Gclc, rotenone significantly reduced total glutathione (**Fig. 3G**) and GSH (**Fig. 3H**) levels. Surprisingly, GSH/GSSG ratio was slightly increased following 4 h of rotenone treatment (**Fig. 3I**). Importantly, pretreatment with N-acetylcysteine (NAC), a precursor that replenishes intracellular glutathione pool, significantly attenuated iron-induced ferroptosis in a dose-dependent manner (**Fig. 3J**). However, NAC supplementation, even at the high dose of 5 mM, failed to rescue RSL3-induced ferroptosis.

Together, our data suggest that acute iron overload induces rapid GSH depletion, followed by mtROS-independent induction of *de novo* glutathione biosynthesis, aimed to limiting oxidative damage and slowing the progression of ferroptosis. In contrast, signaling driven by excessive mtROS production exerts an opposing effect by suppressing glutathione biosynthesis.

### 3.4 mtROS/LP-associated signaling inhibits glycolysis, protecting against ferroptosis

Changes in glucose metabolism, including glycolysis and the pentose phosphate pathway (PPP) (**Fig. 4A**), potentially modulates redox homeostasis and ferroptosis sensitivity. Our RNA-seq data showed that several glycolytic genes were upregulated in response to iron overload. These include genes encoding glucose transporter GLUT1 (SLC2A1), hexokinase 1 and 2 (HK1/2), and pyruvate kinase PKM (**Fig. 4B**). Additionally, genes encoding G6PD and PGD, key enzymes in the PPP, were also significantly upregulated following FAC treatment (**Fig. 4B**). In contrast, the expression of genes encoding glucose transporter GLUT2 (SLC2A2, the primary glucose transporter in hepatocytes ^50^), glucokinase GCK (the predominant hepatic hexokinase ^51^), and aldolase B (ALDOB), was significantly decreased under iron-loaded conditions (**Fig. 4B**). These transcriptional changes in Slc2a2, Gck, and Hk1 were further validated by qRT-PCR (**Fig. 4C**). Notably, the decreased expression of Glut2 and Gck persisted after 24 h of FAC treatment (**Fig. S2**). Alterations in these genes were abolished by pretreatment with either MT or Lip-1 (**Fig. 4B**), suggesting that iron-induced changes in glycolytic gene expression is dependent on mtROS/LP-associated signaling. Interestingly, Lip-1 pretreatment even reversed the direction of gene expression changes relative to iron treatment alone, possibly due to Lip-1 induced reductive stress. Moreover, genes involved in gluconeogenesis, including Pck1, Fbp1 and G6pc, were also decreased by iron overload, though these changes were less dependent on mtROS/LP-associated signaling (**Fig. 4B**).

**Figure 4:**
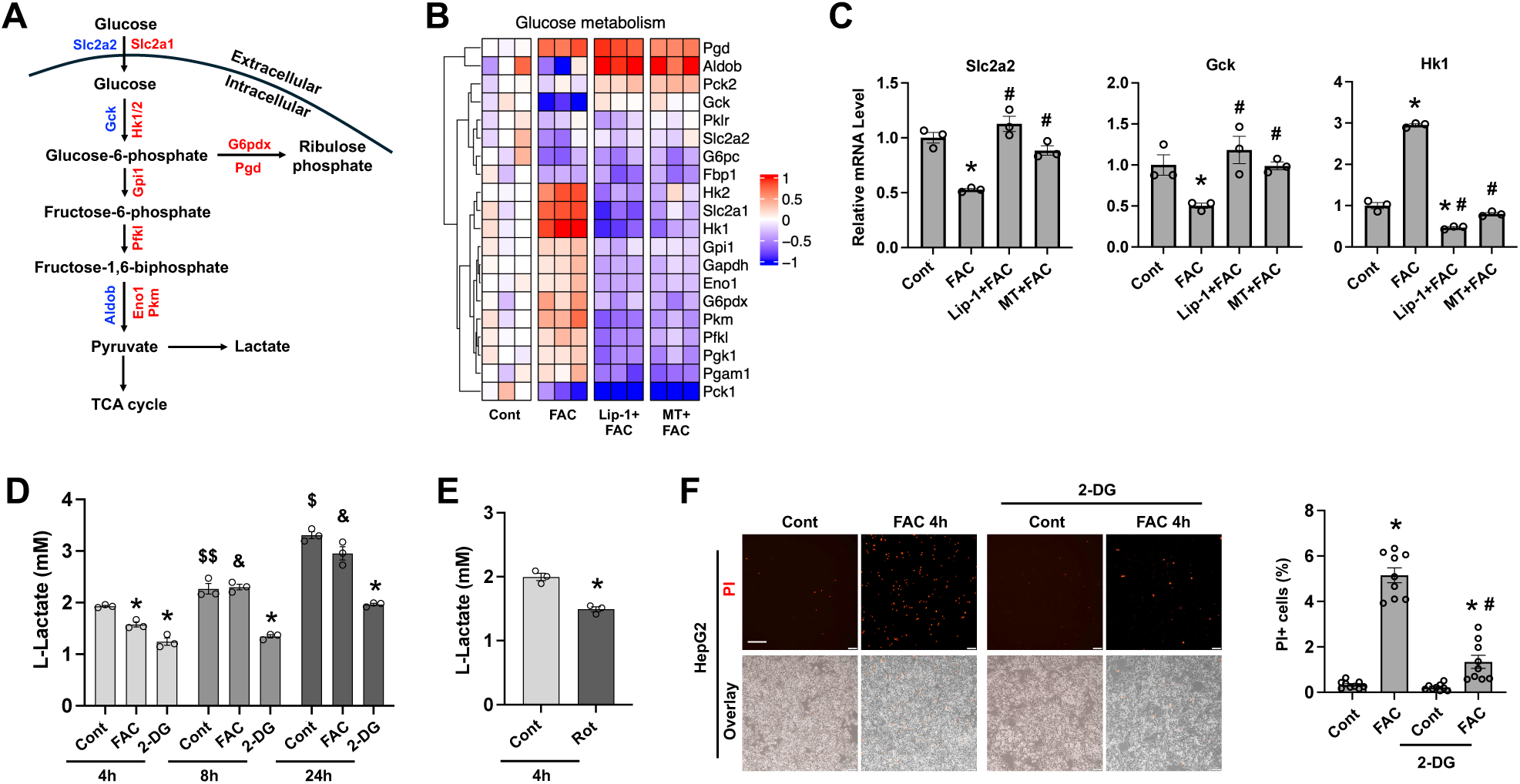
mtROS/LP-dependent downregulation of glycolysis protects against ferroptosis. **(A)** Pathway depicting upregulated (*red*) and downregulated (*blue*) genes involved in glycolysis and pentose phosphate pathway in FAC-treated PMH. (**B**) Heatmap showing the log_2_ fold changes in mRNA expression of genes involved in glucose metabolism. (**C**) qRT-PCR analysis of mRNA expression levels of Slc2a2, Gck, and Hk1. (**D**) L-Lactate concentration in the culture medium of PMH treated with or without 100 μM FAC or 5 mM 2-Deoxy-D-Glucose (2-DG) for the indicated times. (**E**) L-Lactate concentration in the culture medium of PMH treated with or without 200 nM rotenone (Rot) for 4 h. (**F**) HepG2 cells were pretreated for 4 h with or without 5 mM 2-DG, followed by treatment for 4 h with or not 100 μM FAC. Data are mean±SEM. *, *P* < 0.01 compared to Cont within the same group; ^#^, *P* < 0.01 compared to FAC treatment alone; ^$^, *P* < 0.01 and ^$$^, *P* < 0.05 compared to Cont at 4 h; ^&^, *P* < 0.01 compared to FAC-treated cells at 4 h. *Bar*, 200 μm.

The observed mixed expression patterns of glycolytic genes prompted us to assess the functional impact of iron overload on glycolytic activity. PMH were treated with FAC or (2-DG), a glucose analog that inhibits glycolysis ^52^, for different time points. Compared to untreated control, lactate production was significantly decreased in cells treated with FAC for 4h (**Fig. 4D**). This difference, however, was not seen at 8 and 24 h post FAC treatment. Although lactate levels increased at later time points in FAC-treated groups, similar trends were also observed in untreated and 2-DG treated groups (**Fig. 4D**), likely due to the conversion of medium-contained pyruvate to lactate. Our RNA-seq data suggest that mtROS signaling mediates iron-induced glycolytic inhibition. Consistently, rotenone treatment for 4 h also led to a significant reduction in lactate production (**Fig. 4E**), reinforcing the inhibitory effects of mtROS on glycolysis. Our findings indicate that iron excess induces mtROS/LP-dependent suppression of glycolysis in hepatocytes. We then assessed the effects of suppressed glycolysis on ferroptosis susceptibility in hepatocytes and found that pretreatment of HepG2 cells for 4 h with 2-DG reduced the number of PI+ cells following exposure to FAC (**Fig. 4F**). Collectively, these findings suggest that mtROS/LP-initiated signaling suppresses glycolic activity to mitigating further ferroptotic cell death of the hepatocytes.

### 3.5 mtROS/LP-associated signaling impairs fatty acid oxidation, protecting against ferroptosis

LP, particularly the oxidation of polyunsaturated fatty acids (PUFAs), is the critical step in the execution of ferroptosis ^53^. Both fatty acid oxidation and *de novo* lipogenesis of the lipid metabolism rely on proper mitochondrial function ^54^. Our RNA-seq results showed that mRNA expression of Cd36 and fatty acid transport protein 2 (Fatp2/Slc27a2) (**Fig. 5A**), two major fatty acid transporters in hepatocytes ^55,56^, was significantly downregulated in iron-loaded hepatocytes. In contrast, the expression of Fatp1 (Slc27a1) was upregulated, while Fatp4 (Slc27a4) and Fatp5 (Slc27a5) levels remained unchanged (**Fig. 5A**). Most genes involved in the β-oxidation pathway was repressed by FAC treatment, including the gene encoding carnitine palmitoyltransferase 1A (CPT1A) (**Fig. 5A**), a rate-limiting enzyme in fatty acid oxidation ^57^. Pretreatment with MT or Lip-1 prevented iron-induced downregulation of genes involved in fatty acid uptake and oxidation (**Fig. 5A**). The changes in Cd36, Fatp2 and Cpt1a expression was further validated by qRT-PCR analysis (**Fig. 5B**). Moreover, rotenone and antimycin A both induced a marked downregulation of Cd36 and Cpt1a (**Fig. 5C**). Of note, although Cd36 and Cpt1a expression was suppressed in the early phase of treatment by FAC or mtROS inducers, their expression was restored following 24 h of FAC treatment (**Fig. S2**).

**Figure 5:**
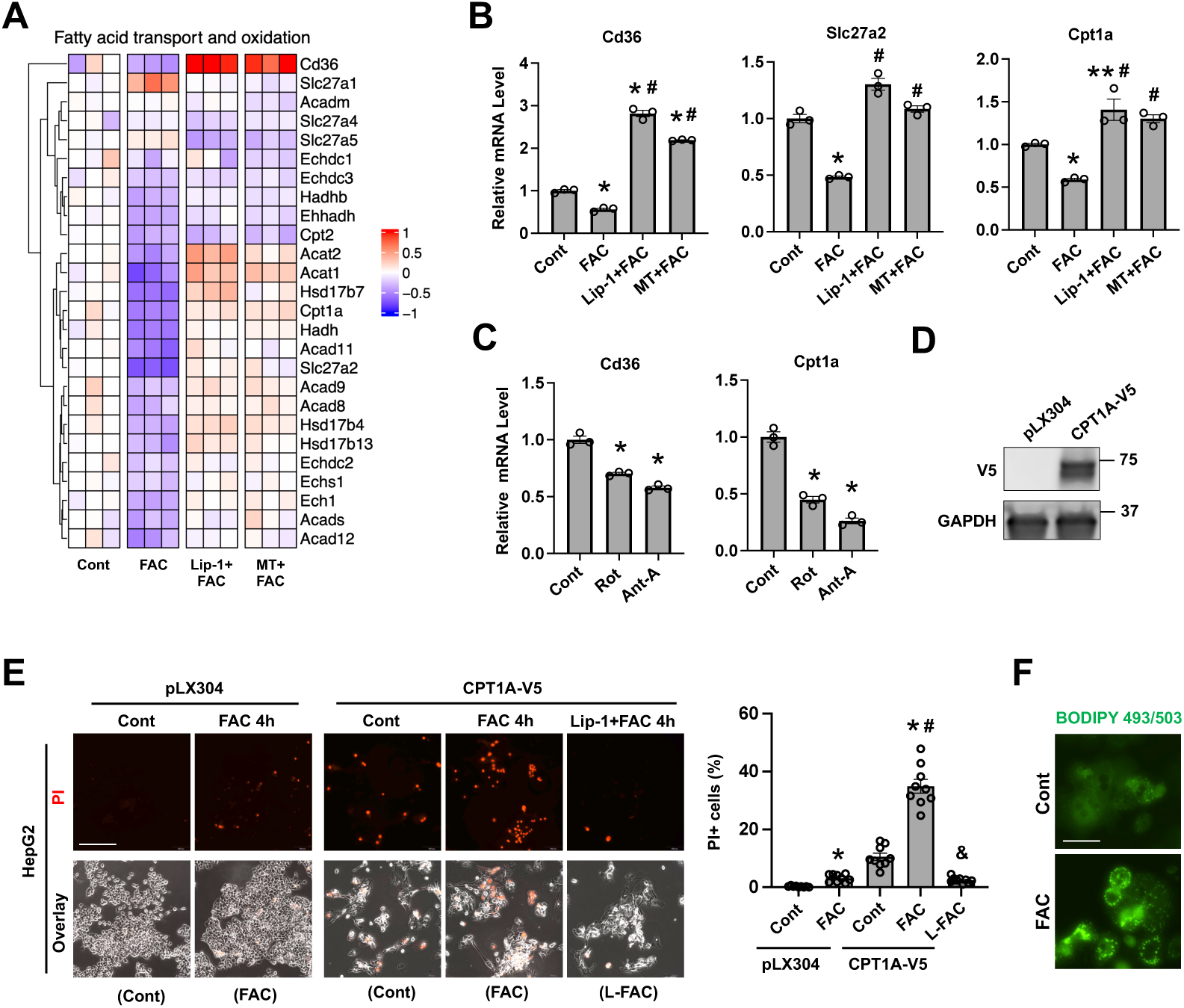
mtROS/LP-dependent downregulation of fatty acid oxidation protects against ferroptosis. (**A**) Heatmap showing the log_2_ fold changes in mRNA expression of genes responsible for fatty acid transport and oxidation. (**B**) qRT-PCR analysis of mRNA expression levels of Cd36, Slc27a2, and Cpt1a. (**C**) Cd36 and Cpt1a mRNA expression in PMH treated for 4 h with or without 200 nM rotenone (Rot) or 10 μM antimycin A (Anti-A). (**D**) Lentivirus-infected HepG2 cells stably expressing CPT1A-V5 or the pLX304 backbone was confirmed by Western blotting with an anti-V5 antibody. (**E**) HepG2/CPT1A-V5 or the pLX304 control cells were pretreated with or without 5 μM Lip-1 prior to treatment with 100 μM FAC for 4 h. Necrotic cells were stained with PI, and the percentage of PI+ cells are shown as mean±SEM (n = 9 fields of 3 independent experiments). (**F**) PMH treated for 24 h with or not 100 μM FAC were stained with BODIPY 493/503. *, *P* < 0.01 and **, *P* < 0.05 compared to Cont within the same group; ^#^, *P* < 0.01 compared to FAC treatment alone (**C**) or FAC-treated Cont cells (**E**); ^&^, *P* < 0.01 compared to FAC-treated CPT1A-V5 cells. *Bars*, 200 μm for *panel* **E** and 50 μm *panel* **F**.

It is generally believed that increased fatty acid oxidation protects against ferroptosis through depletion of the PUFA pool. Herein, we observed that HepG2 cells overexpressing CPT1A were sensitized to iron-induced cell death, and this effect was prevented by Lip-1 pretreatment (**Fig. 5D**). This result suggests that enhancing CPT1A-dependent fatty acid oxidation increases hepatocyte susceptibility to ferroptosis. Interestingly, CPT1A overexpression also promoted PI+ cell death even under basal conditions (**Fig. 5D**). The underlying mechanism is not determined in this study, but it could be related to excessive mtROS production due to hyperactive mitochondrial activity. Together, we speculate that iron overload-induced downregulation of fatty acid uptake and oxidation likely represents a protective mechanism to mitigate oxidative damage and limit ferroptosis of the hepatocytes.

In contrast to the fatty acid uptake and oxidation pathways, genes involved in *de novo* lipogenesis were differentially regulated in iron overload state (**Fig. S4**). Although the specific changes in lipid species remain to be determined in future studies, neutral lipid staining revealed increased accumulation of lipid droplets in iron-loaded hepatocytes (**Fig. 5E**). The Ascl gene family, including Acsl1, Acsl3, and Acsl4, plays critical roles in regulating ferroptosis sensitivity. ACSL1 and ACSL4 promote, whereas ACSL3 protects against ferroptosis, *via* their respective roles in incorporating (PUFAs) versus monounsaturated fatty acids (MUFAs) into the plasma membrane ^58^. Our RNA-seq analysis showed that mRNA expression of Acsl1, the predominant isoform in hepatocytes, was downregulated, whereas Acsl3 expression was upregulated, following acute iron exposure (**Fig. S4**). These changes in Acsl1 and Acsl3 were at least partially dependent on mtROS/LP-associated signaling. In contrast, the expression of Acsl4, a well-characterized positive regulator of ferroptosis ^59^, was not significantly altered in iron-loaded condition.

### 3.6 mtROS/LP-associated signaling suppresses the TCA cycle and mitochondrial respiration

The TCA cycle is a central metabolic pathway that utilizes acetyl-CoA to generate energy precursors NADH and FADH_2_. The above findings of iron-induced mtROS-dependent suppression of glycolysis and fatty acid oxidation leads to reduction in acetyl-CoA production. Additional data from the RNA-seq analysis revealed that the mRNA expression of Idh3a, which encodes the rate-limiting enzyme in the TCA cycle ^60^, was significantly decreased by iron overload (**Fig. 6A**). This downregulation was dependent on mtROS/LP-mediated signaling, as pretreatment with MT or Lip-1 abolished the iron-induced suppression of Idh3a. In contrast, the mRNA expression of Pdha1, which encodes E1-α subunit of the pyruvate dehydrogenase linking glycolysis to the TCA cycle ^61^, was not altered following iron treatment. These transcriptional changes were validated by qRT-PCR analysis (**Fig. 6B**). Moreover, rotenone treatment resulted in the downregulation of Idh3a, further supporting an inhibitory role of the mtROS signaling in regulating TCA reactions (**Fig. 6C**). The TCA cycle requires NAD^+^ as an electron acceptor to produce NADH. We observed significant downregulation of many genes involved in *de novo* NAD^+^ biosynthesis pathway (Tdo2, Ido2, Kmo, Qprt, and Kynu) as well as the NAD+ salvage pathway including (Nmnat1, Nmnat3, and Naprt) in response to iron overload (**Fig. 6D**). Additionally, the expression of Slc25a51, which encodes the mitochondrial NAD^+^ importer, was also significantly reduced in iron-loaded hepatocytes (**Fig. 6D**). These iron-induced changes related to the TCA cycle were dependent on mtROS/LP-driven signaling. By utilizing HepG2 cells stably expressing PDHA1-V5 (**Fig. 6E**), we showed that heightened TCA activity significantly enhanced the sensitivity of hepatocytes to iron-induced ferroptosis (**Fig. 6F**), which is consistent with a previous report that inhibition of pyruvate dehydrogenase attenuates ferroptosis in pancreatic carcinoma cells ^31^.

**Figure 6:**
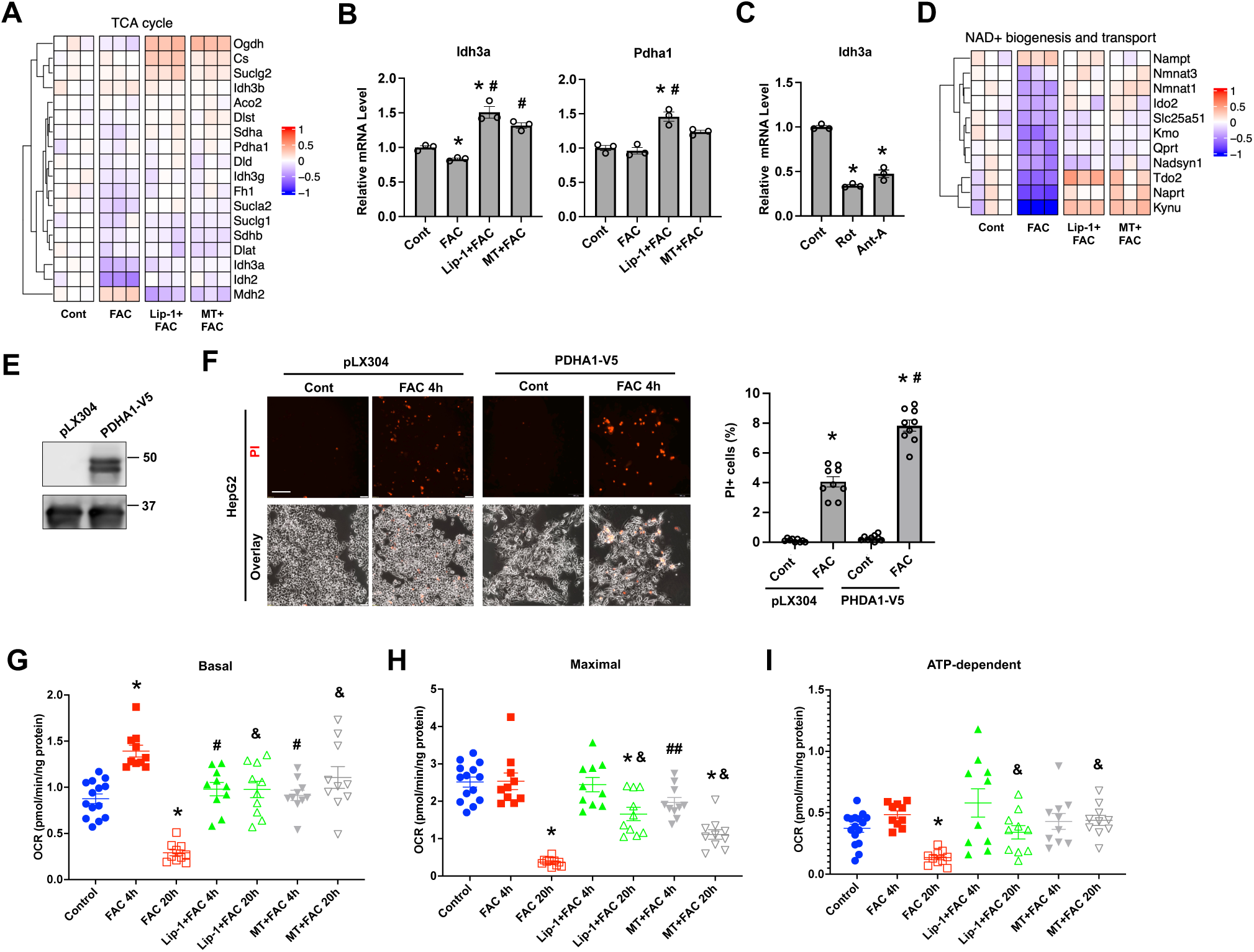
mtROS/LP-associated signaling suppresses TCA cycle activity and mitochondrial respiration. (**A**) Heatmap showing the log_2_ fold changes in mRNA expression of genes of the TCA cycle. (**B**) qRT-PCR analysis of mRNA expression levels of Idh3a and Pdha1. (**C**) Idh3a mRNA expression in PMH treated for 4 h with or without 200 nM rotenone (Rot) or 10 μM antimycin A (Anti-A). (**D**) Heatmap showing the log_2_ fold changes in genes involved in the biogenesis and transport of NAD+. (**E**) PDHA1-V5 expression in HepG2 cells was determined by blotting with an anti-V5 antibody. (**F**) HepG2 cells overexpressing PDHA1-V5 or the pLX304 backbone were treated for 4 h with 100 μM FAC, followed by staining with PI. The percentage of PI+ cells are shown as mean±SEM (n = 9 fields of 3 independent experiments). Basal (**G**), maximal (**H**), and ATP-dependent (**I**) oxygen consumption rate (OCR) was determined in PMH treated with or without MT, Lip-1, or FAC for the indicated times. *, *P* < 0.01 compared to Cont within the same group; ^#^, *P* < 0.01 and ^##^, *P* < 0.05 compared to FAC treatment alone (**B**), FAC-treated pLX304 cells (**F**), and 4-h FAC treatment alone (**G** and **H**); and ^&^, *P* < 0.01 compared to 20-h FAC treatment alone (**G**-**I**). *Bar*, 100 μm.

In line with decreased glycolysis, fatty acid oxidation, and TCA cycle activity, PMH treated with FAC for 24 h showed significant decreases in basal (**Fig. 6G**), maximal (**Fig. 6H**), and ATP-dependent oxygen consumption rates (OCR) (**Fig. 6I**). Pretreatment with MT and Lip-1 prevented FAC-induced decline in basal OCR and improved both maximal and ATP-dependent OCR. Of note, PMH exhibited a significant increase in basal OCR in the early phase (4 h) of FAC treatment (**Fig. 6G**), likely due to an increase in oxygen consumption during iron-catalyzed ROS production.

### 3.7 mtROS/LP-dependent suppression of CoQ biosynthesis enhances ferroptosis sensitivity

CoQ carries electrons from Complex I/II to III in the mitochondrial respiratory chain. Upon accepting electrons, CoQ is reduced to CoQH_2_, a potent antioxidant that protects against LP and ferroptosis ^14–16^. CoQ biosynthesis involves the synthesis of a benzoquinone head group and a polyisoprenoid tail in the cytoplasm, followed by their assembly and modification within the inner mitochondrial membrane via the CoQ biosynthetic complex ^62^. The synthesis of is dependent on the mevalonate pathway ^63^. Our RNA-seq analysis identified that iron overload significantly downregulated the expression of many genes involved in the mevalonate and cholesterol biosynthesis pathways (**Fig. 7A**). Notably, this iron-induced suppression of the mevalonate pathway was reversed by pretreatment with MT or Lip-1. The transcription factor Sterol Regulatory Element Binding Protein 2 (SREBP-2) is master regulator of the mevalonate pathway ^64^, and is known to be inhibited by the AMPK signaling ^65^. We showed that mtROS induction by rotenone activated AMPKα and reduced nuclear SREBP-2 levels (**Fig. 7B**), suggesting mtROS is an important upstream regulator of the AMPK-SREBP-2 axis. Moreover, treatment with RSL3, which induces excessive mtROS production ^24^, also led to significantly reduced expression of nuclear form SREBP-2 (**Fig. S5A**). In contrast, scavenging lipid ROS by Lip-1 at the basal condition did not modulate SREBP-2 expression (**Fig. S5A**). Tyrosine metabolism is the primary source of 4-hydroxybenzoate (4-HB), a precursor for the benzoquinone head group of CoQ ^62^. We observed that the expression of Tat, which encodes the rate-limiting enzyme tyrosine aminotransferase mediating tyrosine degradation in the liver ^66^, was significantly reduced in iron-loaded hepatocytes (**Fig. 7C**).

**Figure 7:**
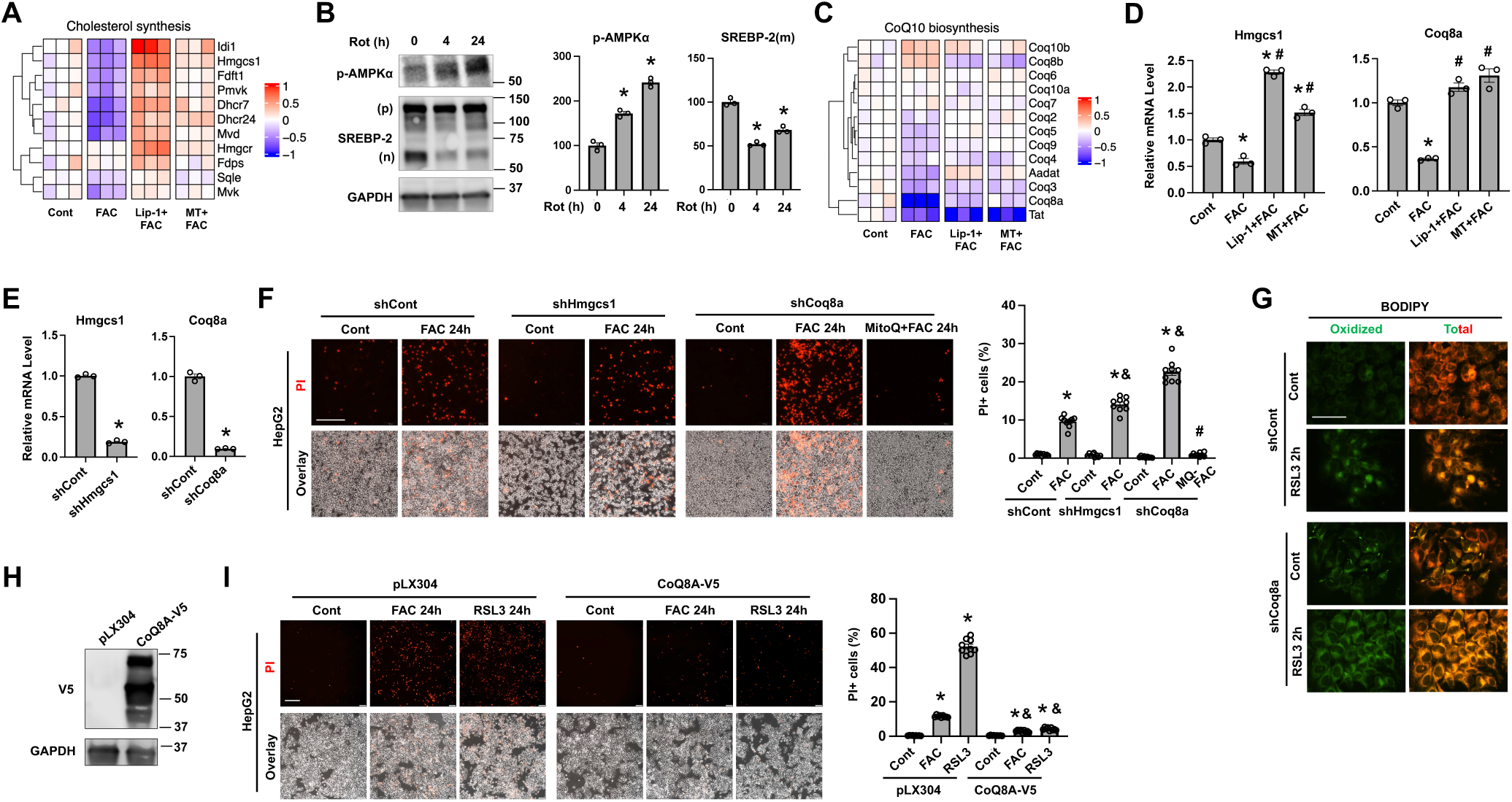
mtROS/LP-dependent suppression of CoQ biosynthesis sensitizes hepatocytes to ferroptosis. (**A**) Heatmap showing the log_2_ fold changes in mRNA expression of genes in the mevalonate/cholesterol synthesis pathway. (**B**) Changes in the protein expression of phospho-AMPKα, precursor (p) and nuclear (n) forms of SREBP-2 in HepG2 cells treated for the indicated times with 200 nM rotenone. (**C**) Heatmap showing the log_2_ fold changes in mRNA expression of genes responsible for CoQ biosynthesis. (**D**) qRT-PCR analysis of mRNA expression levels of Hmgcs1 and Coq8a. (**E**) qRT-PCR analysis of Hmgcs1 and Coq8a mRNA expression in HepG2 cells with lentiviral knockdown. (**F**) HepG2 cells with the knockdown of HMGCS1 (shHmgcs1), CoQ8A (shCoq8a), or scrambled control (shCont) were treated for 24 h with 100 μM FAC, followed by staining with PI. (**G**) BODIPY 493/503 staining of RSL3-treated HepG2 cells with or without the knockdown of CoQ8A. (**H**) CoQ8A-V5 expression in HepG2 cells was determined by Western blotting with an anti-V5 antibody. (**I**) HepG2 cells expressing CoQ8A-V5 or pLX304 backbone were treated for 24 h with 100 μM FAC or 100 nM RSL3, followed by staining with PI. Data are shown as mean±SEM.*, *P* < 0.01 compared to Cont within the same group; ^#^, *P* < 0.01 compared to FAC treatment alone; and ^&^, *P* < 0.01 compared to FAC treatment in shCont (**F**) and pLX304 (**H**) cells. *Bars*, 200 μm for *panel*s **F** and **I,** and 50 μm *panel* **G**.

The CoQ biosynthetic complex, located at the inner mitochondrial membrane, consists of many proteins including CoQ1-10, of which CoQ8A plays a critical role in complex assembly by phosphorylating CoQ3, CoQ5, and CoQ7 ^67,68^. Interestingly, Coq8a mRNA expression in PMH was markedly downregulated following iron treatment (**Fig. 7C**). Our qRT-PCR analysis further confirmed iron-induced mtROS/LP-dependent suppression of Coq8a and Hmgcs1 expression (**Fig. 7D**). Moreover, persistent Coq8a downregulation was observed following 24 h of FAC treatment (**Fig. S2**). The Coq8a expression was also reduced by RSL3 treatment (**Fig. S5B**).

Hmgcs1 and Coq8a are both required for CoQ biosynthesis. Thus, we determined their roles in regulating the sensitivity of hepatocytes to ferroptosis by using HepG2 cells with silenced expression of HMGCS1 or CoQ8A (**Fig. 7E**). In this study, only the mRNA expression of Coq8a was determined because we were unable to obtain a reliable antibody against CoQ8A protein. HMGCS1 knockdown significantly but only modestly increased the sensitivity of HepG2 cells to FAC-induced ferroptosis compared to the control. In contrast, CoQ8A knockdown led to a marked increase in the number of PI+ cells in response to FAC treatment (**Fig. 7F**). Notably, the increase was abolished by pretreatment with MitoQ (**Fig. 7F**), a mitochondria-targeted analog of ubiquinol. Consistently, CoQ8A knockdown led to increased LP at both basal and RSL3-treated conditions (**Fig. 7G**). The pro-ferroptotic effects of CoQ8A deficiency were also observed in multiple cancer cell lines, including MIA Paca-2, H1299, and MDA-MB-231 cells, although different cell types exhibited varying sensitivities to iron overload and RSL3 treatment (**Fig. S6**). Using HepG2 cells overexpressing CoQ8A-V5 (**Fig. 7H**), we further showed that elevated CoQ8A expression conferred strong protection against ferroptosis induced by FAC and RSL3 (**Fig. 7I**).

Our findings suggest that mtROS/LP-driven signaling downregulates CoQ biosynthesis by disrupting the mevalonate pathway and CoQ8A-mediated assembly of the CoQ synthetic machinery. These changes were likely adopted by the hepatocytes to limit mitochondrial respiration under oxidative stress. However, the resulting CoQ deficiency increases hepatocyte susceptibility to ferroptosis, particularly under conditions of cystine or glutathione deprivation, or when GPx4 activity is compromised.

### 3.8 mtROS/LP induces FXR/RXRα-dependent downregulation of CoQ8A expression

We next sought to elucidate the mechanism underlying the transcriptional downregulation of Coq8a. Our RNA-seq data showed that several nuclear receptors involved in lipid metabolism displayed expression patterns mimicking Coq8a in response to iron overload, MT, and Lip-1 treatments. These include retinoic X receptor (RXR) family members (RXRα, β, and γ), farnesoid X receptor (FXR/NR1H4), and liver X receptor (LXR/NR1H3) (**Fig. 8A**). In contrast, the mRNA expression of peroxisome proliferator-activated receptor PPARγ was increased by iron overload, whereas PPARα expression was unchanged. Immunoblotting analysis further confirmed a significant decrease in RXRα protein expression in PMH treated with FAC (**Fig. 8B**), rotenone (**Fig. 8C**), or RSL3 (**Fig. S5C**) for 4 h and 24 h. We also observed a significant decrease in Fxr expression in response to rotenone treatment (**Fig. 8D**). To evaluate the roles of these nuclear receptors in Coq8a regulation, we first utilized a panel of pharmacological agonists and antagonists, including GW7647 (PPARα agonist), GW6471 (PPARα antagonist), bexarotene (RXR agonist), HX531 (RXR antagonist), LXR623 (LXR agonist), GSK2033 (LXR antagonist), and fexaramine (FXR agonist). We identified that Coq8a transcript level was significantly decreased by HX531 and increased by fexaramine (**Fig. S7**), implicating RXR and FXR as potential positive regulators of Coq8a gene transcription. Activation of RXR alone (via bexarotene) did not affect Coq8a expression, likely because RXR typically functions as a heterodimer with other nuclear receptors ^69^. Consistent with Coq8a, the mRNA expression of Rxra and Fxr remained low after 24 h of FAC treatment (**Fig. S2**). APAP overdose induces acute liver failure in part through increasing mtROS production and mitochondrial dysfunction. We found that RXRα protein and Coq8a mRNA expression were also significantly decreased *in vivo* in livers of APAP-treated mice (**Fig. S8**).

**Figure 8:**
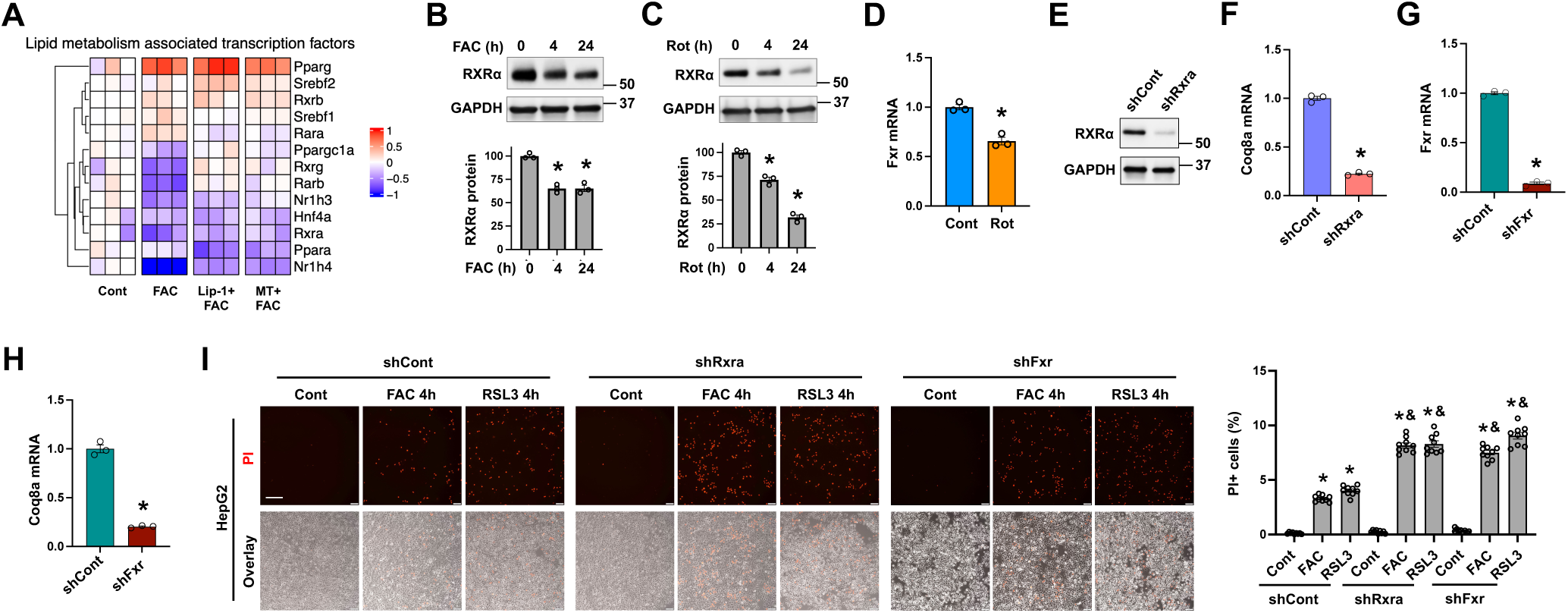
FXR/RXRα regulates CoQ8A expression and ferroptosis sensitivity of hepatocytes. (**A**) Heatmap showing the log_2_ fold changes in mRNA expression of transcription factors that regulate hepatic lipid metabolism. RXRα protein expression in PMH treated for the indicated times with 100 μM FAC (**B**) or 200 nM rotenone (Rot) (**C**). (**D**) Fxr mRNA level in PMH treated for 4 h with or without 200 nM Rot. (**E**) RXRα expression in HepG2 cells with stable knockdown of RXRα and control cells. (**F**) Coq8a mRNA expression in HepG2 cells with or not the knockdown of RXRα. (**G**) Fxr mRNA expression in HepG2 cells with stable knockdown of FXR and control cells. (**H**) Coq8a mRNA expression in HepG2 cells with or not the knockdown of FXR. (**I**) HepG2 cells with or without the knockdown of RXRα or FXR were treated for 4 h with 100 μM FAC or 100 nM RSL3, followed by staining with PI. Data are mean±SEM.*, *P* < 0.01 compared to Cont within the same group; and ^&^, *P* < 0.01 compared to FAC or RSL3 treatment of shCont cells. *Bar*, 200 μm.

To confirm the role of RXR in Coq8a regulation, we generated HepG2 cells with the knockdown of RXRα (**Fig. 8E**), the most abundant isoform of RXR in hepatocytes ^70^. Knockdown of RXRα led to a significant reduction in Coq8a mRNA levels (**Fig. 8F**). Similarly, FXR knockdown resulted in an approximately 80% decrease in Coq8a expression compared to the control cells (**Fig. 8G-H**). Next, we assessed how deficiencies in RXRα and FXR modulate ferroptosis sensitivity of the hepatocytes. Compared to control cells, knockdown of either RXRα or FXR significantly increased the number of PI^+^ cells following FAC or RSL3 treatment (**Fig. 8I**). These effects are independent of GPx4 or FSP1, because their expression remained unchanged in both knockdown conditions (**Fig. S9**). Collectively, our data demonstrate that RXRα and FXR cooperatively regulate Coq8a gene transcription, playing an important role in modulating ferroptosis sensitivity of the hepatocytes.

## 4. Discussion

Our study identifies that mtROS induces adaptive and maladaptive changes in cellular metabolism and antioxidant defenses, exerting distinct effects on hepatocyte susceptibility to ferroptosis. On one hand, mtROS-associated signaling suppresses glycolysis, fatty acid oxidation, and TCA cycle activity, collectively protecting hepatocytes against ferroptosis (**Fig. 9**). On the other hand, mtROS-driven signaling impairs glutathione biosynthesis by reducing cystine uptake and diminishing the abundance of glutamate-cysteine ligase. In parallel, mtROS downregulates the mevalonate pathway and CoQ8A expression, leading to suppressed CoQ biosynthesis (**Fig. 9**). Impaired glutathione and CoQ production, together with elevated ROS production, results in the depletion of their reduced forms - GSH and CoQH_2_, two key antioxidants that defend cells against LP-dependent ferroptosis. Our RNA-seq analysis further demonstrates that these mtROS-induced changes are largely dependent on activation of LP-driven signaling.

**Figure 9:**
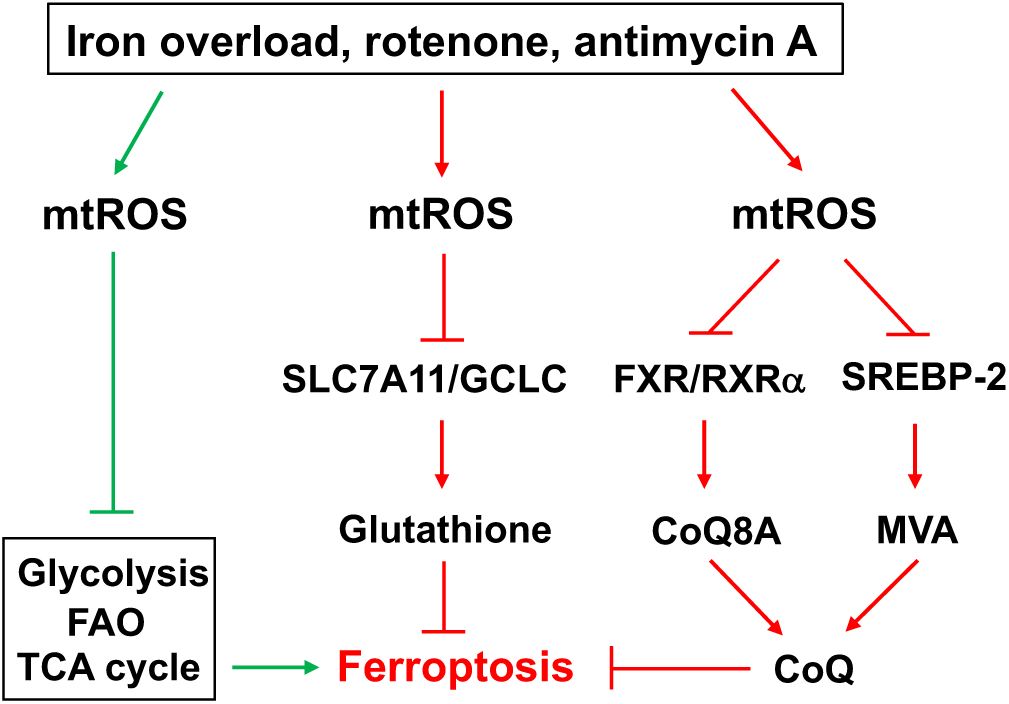
A summary of mtROS-dependent metabolic alterations that differentially regulate ferroptosis sensitivity of the hepatocytes. The mtROS-associated signaling downregulates glycolysis, fatty acid oxidation (FAO), and tricarboxylic acid (TCA) cycle activity, protecting hepatocytes from undergoing ferroptosis. However, excessive mtROS suppresses the biosynthesis of glutathione and coenzyme Q10 (CoQ), increasing hepatocyte susceptibility to ferroptosis. The mtROS-induced signaling inhibits glutathione biosynthesis by downregulating SLC7A11 and GCLC, and reduces CoQ biosynthesis through inhibition of SREBP-2-dependent mevalonate (MVA) pathway and FXR/RXRα-mediated CoQ8A expression.

A substantial body of work has shown that alterations in glycolysis, fatty acid oxidation, and TCA cycle activity modulate cellular sensitivity to ferroptosis. Impaired glycolysis or fatty acid oxidaiton can either enhance or reduce vulnerability of cells to ferroptosis depending on the contexts. The PPP, which branches with the early steps of glycolysis (glucose uptake and glucose-6-phosphate production), contributes to NADPH production to support the regeneration of GSH and CoQH_2_ ^71^. Conversely, enhanced glycolic flux increases the availability of pyruvate, activating the TCA cycle and *de novo* synthesis of PUFAs ^31^, thereby promoting ferroptosis. Similarly, increased fatty acid oxidation has been proposed to confer protection against ferroptosis by reducing PUFA levels ^72^. Other studies, however, suggest that fatty acid oxidation increases acetyl-CoA availability fueling the TCA cycle, leading to increased ROS production ^73–75^. In contrast to the context-dependent roles of glycolysis and fatty acid oxidation in modulating ferroptosis sensitivity, the impact of TCA cycle activity on ferroptosis sensitivity is relatively consistent. It was demonstrated that enhancing TCA reaction (e.g., glutaminolysis) increases ferroptosis vulnerability ^29^. Conversely, impaired TCA cycle activity due to inhibition of the pyruvate dehydrogenase (PDH) by pyruvate dehydrogenase kinase 4 leads to ferroptosis resistance ^31^. In this study, we demonstrate that mtROS-dependent signaling downregulates glycolysis, fatty acid oxidaiton, and TCA cycle activity. Iron overload and pharmacological mtROS inducers not only reduce glucose and fatty acid uptake but also suppress the expression of rate-limiting enzymes in glycolysis and β-oxidation. Moreover, mtROS signaling inhibits TCA cycle activity by diminishing the availability of NAD+ and downregulating IDH3A – an enzyme that catalyzes the rate-limiting step of the TCA cycle by irreversibly converting isocitrate to alpha-ketoglutarate ^76^. Importantly, our current study showed that acute inhibition (4-h treatment by 2-DG) of glycolysis mitigated ferroptosis, whereas upregulation of fatty acid oxidation (by overexpressing CPT1A) and TCA cycle activity (by overexpressing PDHA1) promoted ferroptosis, of cultured hepatocytes, at least in the scenario of iron overload induced ferroptosis. Therefore, suppression of glycolysis, fatty acid oxidaiton and TCA reactions by mtROS-driven signaling are likely adaptive responses that hepatocytes develop to limit further oxidative damage and the rate of ferroptosis. This interpretation is supported by our previous observation that hepatocytes with increased availability of glucose, fructose, fatty acids, and amino acids exhibit heightened vulnerability to ferroptosis ^24^. However, we do not exclude the possibility that prolonged downregulation of glycolysis, whether caused by glucose deprivation or reduced expression of glucokinases, may increase the susceptibility of hepatocytes to ferroptosis due to insufficient production of NADPH.

GSH provides the primary line of defense against ROS-induced cellular damage ^77^. Our findings reveal that GSH biosynthesis is differentially regulated under conditions of iron overload and pharmacological mtROS induction. Iron overload enhances glutathione biosynthesis by increasing cellular uptake of precursor amino acids and upregulating the expression of GCLC and GCLM, which catalyze glutathione biosynthesis. In contrast, treatment with rotenone or antimycin A reduces the expression of SLC7A11 and GCLC and glutathione levels, consistent with previous reports of the effects of mtROS inducers ^78–80^. It is worth noting that, although acute iron overload promotes mtROS production, it also catalyzes ROS production in extra-mitochondrial compartments ^25^. Supporting a role for non-mitochondrial ROS associated signaling in stimulating glutathione biosynthesis, our data show that scavenging mtROS or lipid ROS in iron-loaded hepatocytes does not abolish but rather enhances the expression of genes involved in glutathione biosynthesis. It remains unclear why cellular homeostasis program downregulates *de novo* glutathione biosynthesis under conditions of excessive mtROS.

Coenzyme Q (CoQ) is an essential electron carrier in the mitochondrial electron transport chain, and its reduced form, CoQH₂, functions as a potent lipid-soluble antioxidant ^81^. More recently, CoQH₂ is identified as a potent anti-ferroptosis lipid species, the generation of which is dependent on the activities of FSP1 ^14–16^, DHODH ^18^, mitochondrial complex I and II ^17^. Consequently, impaired CoQH₂ regeneration, or CoQ deficiency resulting from defects in CoQ biosynthesis, sensitizes cells to ferroptosis, particularly when the GPx4–GSH axis is compromised. In this study, we demonstrate that mtROS/LP-driven signaling markedly suppress the expression of multiple genes indispensable for CoQ biosynthesis through both the mevalonate pathway and the CoQ biosynthetic complex. Our study suggests a promoting effect of AMPK on hepatocyte ferroptosis by suppressing the activity of SREBP-2, a master regulator of the mevalonate pathway ^64^. This role of AMPK is in line with a previous report that AMPK activation increases ferroptosis vulnerability by suppressing cystine uptake and glutathione biosynthesis ^82^. Notably, studies have also shown inhibitory effects of AMPK on ferroptosis in cancer cells by limiting the availability of PUFAs ^32,83^, suggesting complex and context-dependent roles of AMPK in regulating ferroptosis sensitivity. Moreover, for the first time, we have identified that mtROS-dependent signaling downregulates CoQ8A, an atypical kinase essential for the assembly of the CoQ biosynthetic complex by phosphorylating and stabilizing CoQ3, CoQ5, and CoQ7 ^67,68^. As such, loss-of-function mutations in Coq8a in humans cause CoQ deficiency and early-onset cerebellar ataxia ^84^. Our work provides the direct evidence that CoQ8A expression inversely modulate the susceptibility of hepatocytes and multiple cancer cell lines to ferroptosis. Interestingly, a recent study has shown that radiotherapy-induced CoQ8A upregulation mediates radioresistance of lung cancer cells ^85^.

Our RNA-seq analysis further revealed that several nuclear receptors involved in hepatic lipid metabolism are downregulated by iron overload in an mtROS/LP-dependent manner, suggesting a potentially important role for the mtROS-nuclear receptor signaling axis in the feedback regulation of lipid metabolism under pathological liver conditions. Specifically, we identified FXR and RXRα as critical regulators of Coq8a expression, and deficiency in FXR or RXRα sensitizes hepatocytes to ferroptosis. Our finding of the protective roles of FXR and RXRα against ferroptosis is consistent with a previous study using fibrosarcoma HT-1080 cells, which showed that FXR and RXRα positively regulate the expression of GPx4 and FSP1 ^86^. However, in hepatocytes, we did not observe similar effects on GPx4 and FSP1 expression. Instead, our findings suggest that FXR and RXRα collaborate to protect against ferroptosis in part through CoQ8A-dependent CoQ biosynthesis. Of note, the mtROS-mediated downregulation of the FXR/RXRα-CoQ8A axis is not only seen in cultured cells but also *in vivo* under conditions of APAP overdose-induced liver injury. It would be interesting to determine in the future whether downregulations of mevalonate pathway and CoQ8A occurs and contributes to liver injury associated with ischemia-reperfusion, which often characterized by excessive mtROS production ^87,88^.

In conclusion, we have identified previously unrecognized mechanisms by which mtROS-driven signaling enhances ferroptosis susceptibility through suppression of glutathione and CoQ biosynthesis. Given that mitochondrial dysfunction and excessive mtROS production are hallmark features of many pathological conditions, including mitochondrial diseases, our findings suggest that mtROS-dependent suppression of CoQ synthesis may exacerbate disease progression by enhancing ferroptosis sensitivity. Moreover, CoQ insufficiency, together with mtROS-induced downregulation of glycolysis, fatty acid oxidation, and TCA cycle activity, can further aggravate metabolic abnormalities. An important remaining question is whether and how distinct levels of mtROS may exert differential effects on cell metabolism.

## Acknowledgements

This study was supported by NIH grant R01DK125647 (to PH) and R01DK140441 (to PH) as well as a VA Career Development Award 5IK2BX005913-02 (to MRS). Next generation sequencing services were provided by the Emory NPRC Genomics Core which is supported in part by NIH P51 OD011132. Sequencing data was acquired on an Illumina NovaSeq 6000 funded by NIH S10 OD026799. We thank Dr. Xiulei Mo (Emory University) for providing the MIA Paca-2 cell line used in this study.

The content is solely the responsibility of the authors and does not necessarily represent the official views of the U.S. Department of Veterans Affairs.

## Author contributions

Conception and design: PH; Acquisition of data: SB, MRS, YIW, MW; Analysis and interpretation of data: SB, MRS, YIW, DG, SK, GKT, PH; Writing, review, and revision of the manuscript: SB, MRS, SK, PH.

## Abbreviations

2-DG: 2-Deoxy-D-Glucose
CoQ: coenzyme Q10
FAC: ferric ammonium citrate
FSP1: ferroptosis suppressor protein 1
FXR: farnesoid X receptor
GPx4: glutathione peroxidase 4
GSH: reduced glutathione
GSSG: oxidized glutathione
Lip-1: liproxstatin-1
LP: lipid peroxidation
MT: Mito-TEMPO
mtROS: mitochondrial ROS
NAC: N-acetylcysteine
PI: propidium iodide
PMH: primary mouse hepatocytes
PPP: pentose phosphate pathway
RXR: retinoid X receptor

