## Supplementary documents for "Mitochondrial ROS-induced metabolic alterations differentially regulate ferroptosis sensitivity"

**Supplementary Table 1. List of reagents, antibodies, and plasmid constructs.**

| Name | Source | Catalog | Species |
|---|---|---|---|
| **Reagents** | | | |
| (1S, 3R)-RSL3 | Cayman | 19288 | |
| 2-Deoxy-D-Glucose (2-DG) | Cayman | 14325 | |
| Acetaminophen | Cayman | 10024 | |
| Antimycin A | Enzo Life Sciences | 380-075-M010 | |
| Bexarotene | Cayman | 11571 | |
| Bicinchoninic Acid Assay kit | Sigma-Aldrich | BCA1 | |
| Blasticidin | Thermo Fisher Scientific | R210-01 | |
| BODIPY 493/503 | Thermo Fisher Scientific | D3922 | |
| Cell Lysis Buffer (10x) | Cell Signaling Technology | 9803 | |
| Halt Protease and Phosphatase Inhibitor | Thermo Fisher Scientific | 78446 | |
| Dexamethasone | Sigma-Aldrich | D2915 | |
| DMEM, high glucose | Thermo Fisher Scientific | 11965092 | |
| Ferric Ammonium Citrate | Sigma-Aldrich | F5879 | |
| Fetal Bovine Serum | R&D Systems | S11150 | |
| Fexaramine | Cayman | 17369 | |
| Glutathione Assay Kit | Cayman | 703002 | |
| Glycolysis Cell-Based Assay Kit | Cayman | 600450 | |
| GSK2033 | Cayman | 25443 | |
| GW6471 | Cayman | 11697 | |
| GW7647 | Cayman | 10008613 | |
| HBSS (10x) | Gibco | 14185-052 | |
| HEPES (1M) | Gibco | 15630-080 | |
| High-Capacity cDNA Reverse Transcription Kit | Applied Biosystems | 43-688-13 | |
| HX531 | Cayman | 20762 | |
| Image-iT Lipid Peroxidation Kit | Thermo Fisher Scientific | C10445 | |
| Insulin | Sigma-Aldrich | I0516 | |
| Liberase | Sigma-Aldrich | 05401119001 | |
| Lipofectamine 3000 | Thermo Fisher Scientific | L3000015 | |
| Liproxstatin-1 | Cayman | 17730 | |
| LXR-623 | Cayman | 21117 | |
| Mito-TEMPO | Cayman | 18796 | |
| N-acetylcysteine (NAC) | MP Biomedicals | 194603 | |
| Percoll (40%) | Sigma-Aldrich | P1644 | |
| Polybrene | Sigma-Aldrich | TR-1003 | |
| *Power* SRBR Green PCR Master Mix | Applied Biosystems | 43-676-59 | |
| Propidium Iodide (PI) | Invitrogen | P1304MP | |
| Puromycin | Sigma-Aldrich | P8833 | |
| RNeasy Mini Kit | Qiagen | 74136 | |
| Rotenone | Cayman | 13995 | |

| | | | |
|---|---|---|---|
| RPMI Medium | Sigma-Aldrich | R8758 | |
| William's E Medium | Thermo Fisher Scientific | 12551032 | |
| William's E Medium (no phenol red) | Thermo Fisher Scientific | A1217601 | |
| **Antibodies** | | | |
| anti-mouse HRP | Cell Signaling Technology | 7076 | Goat |
| anti-rabbit HRP | Cell Signaling Technology | 7074 | Goat |
| FSP1 | Cell Signaling Technology | 24972 | Rabbit |
| GAPDH | Cell Signaling Technology | 2118 | Rabbit |
| GPx4 | Cell Signaling Technology | 59735 | Rabbit |
| Phospho-AMPKα | Cell Signaling Technology | 2535 | Rabbit |
| RXRα | Cell Signaling Technology | 3085 | Rabbit |
| SREBP-2 | Cell Signaling Technology | 25940 | Rabbit |
| V5 | Cell Signaling Technology | 13202 | Rabbit |
| **Plasmid Constructs** | | | |
| pLKO.1/shCoq8a | Sigma-Aldrich | TRCN0000315428 | Human |
| pLKO.1/shNr1h4(Fxr) | Sigma-Aldrich | TRCN0000428528 | Human |
| pLKO.1/shHmgcs1 | Sigma-Aldrich | TRCN0000218729 | Human |
| pLKO.1/shRxra | Sigma-Aldrich | TRCN0000330707 | Human |
| pLenti6.3/COQ8A-V5 | DNASU | HsCD00937726 | Human |
| pLX304/CPT1A-V5 | DNASU | HsCD00444121 | Human |
| pLX304/PDHA1-V5 | DNASU | HsCD00444246 | Human |

**Supplementary Table 2. List of primers and sequences**

| Mouse genes | Sequence | Mouse genes | Sequence |
|---|---|---|---|
| Atp7b-F | CTTCGAAGCGTCAGTCATGG | Lrp8-F | GTGGAAGTAGCCACCAATCGCA |
| Atp7b-R | GGATCGAACTTCACATGGGC | Lrp8-R | CTGCTCATCAATGAGGACCACC |
| Cd36-F | GGACATTGAGATTCTTTTCCTCTG | Pdha1-F | GTGAGAACAACCGCTATGGCATG |
| Cd36-R | GCAAAGGCATTGGCTGGAAGAAC | Pdha1-R | CGCAAACTTTGTTGCCTCTCGG |
| Coq8a-F | CCTCACGGCTGAAGACATTG | Rpl13a-F | ATGACAAGAAAAAGCGGATG |
| Coq8a-R | CCTACAGCCAGACCTCCAAA | Rpl13a-R | CTTTTCTGCCTGTTTCCGTA |
| Cpt1a-F | GGCATAAACGCAGAGCATTCCTG | Slc2a2-F | GTTGGAAGAGGAAGTCAGGGCA |
| Cpt1a-R | CAGTGTCCATCCTCTGAGTAGC | Slc2a2-R | ATCACGGAGACCTTCTGCTCAG |
| Fxr-F | GGGATGAGTGTGAAGCCAGCTA | Slc7a11-F | GGGGAAGGTGATGGCTGTAT |
| Fxr-R | GTGGCTGAACTTGAGGAAACGG | Slc7a11-R | CGCACAAATACCATCGAGCA |
| Gck-F | GCATCTCTGACTTCCTGGACAAG | Slc25a37-F | TCTCACAACCATCACCCTCC |
| Gck-R | CTTGGTCCAGTTGAGCAGGATG | Sc25a37-R | AGCAACACTAGGCCACTCTT |
| Gclc-F | AAGCCTCCTCCTCCAAACTC | Slc27a2-F | TTCAACAGCGGAGACCTCCTGA |
| Gclc-R | GGGCCACTTTCATGTTCTCG | Slc27a2-R | CCACGATGTCAGCGACTTCTGT |
| Gclm-F | TCCTGCTGTGTGATGCCACCAG | Slc30a1-F | CAACACCAGCAATTCCAACGGG |
| Gclm-R | GCTTCCTGGAAACTTGCCTCAG | Slc30a1-R | CGCTTCCAGATTGTCAGACTCC |
| Hk1-F | GAAAGGAGACCAACAGCAGAGC | Slc40a1-F | TGGAATTTGGTGTCCATGTG |
| Hk1-R | TTCGTTCCTCCGAGATCCAAGG | Slc40a1-R | TCAAGTTCACGGATGTTGGA |
| Hmgcs1-F | CAGCTCTTTCACCATGCCTG | Tfrc-F | AATGGGTGTTGGGAAGACAA |
| Hmgcs1-R | ACCACAGTCAGGCAAAGAGA | Tfrc-R | ACATTCTCAGGTGGCAGCTT |
| Idh3a-F | GCAGGACTGATTGGAGGTCTTG | | |
| Idh3a-R | GCCATGTCCTTGCCTGCAATGT | | |
| **Human genes** | **Sequence** | | |
| Coq8a-F | CAACATCCTGGTTCTGTGCC | | |
| Coq8a-R | GGTCGGTGAAGGATCTGTCA | | |
| Fxr-F | ACTTCCGTCTGGGCATTCTGAC | | |
| Fxr-R | GCTGTAAGCAGAGCATACTCCTC | | |
| Hmgcs1-F | TCAGCGAAGACATCTGGTGCCA | | |
| Hmgcs1-R | AAGTCACACAAGATGCTACACCG | | |
| Rpl13a-F | CTCAAGGTGTTTGACGGCATCC | | |
| Rpl13a-R | TACTTCCAGCCAACCTCGTGAG | | |

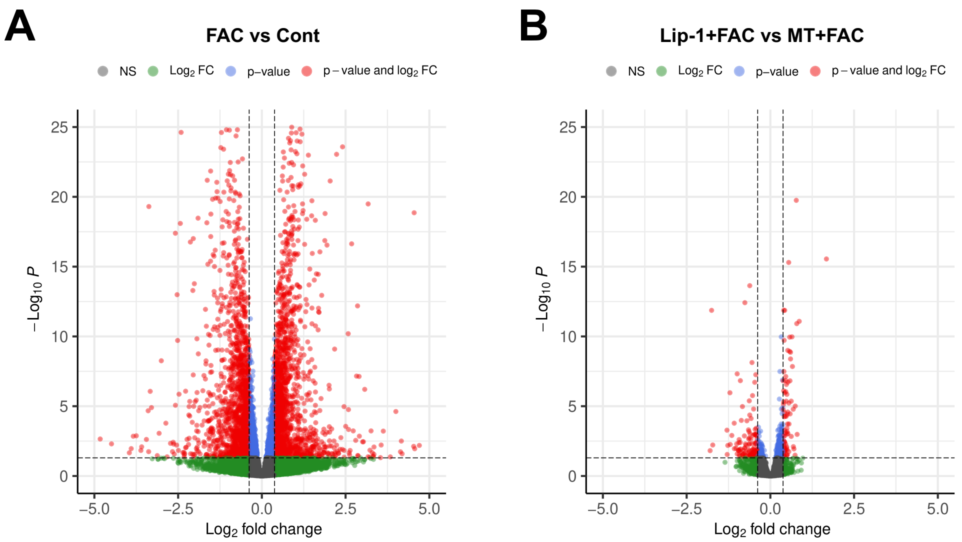

**Figure S1: The effects of iron overload and antioxidant treatment on gene expression profiles in PMH**. Cultured PMH were pretreated with or without 5 μM liproxstatin-1 (Lip-1) or 10 μM Mito-TEMPO (MT) prior to treatment for 4 h with or without 100 μM ferric ammonium citrate (FAC). Cells were harvested for bulk RNA-seq analysis of mRNA expression profiles. (**A**) Volcano plot showing overall changes in the mRNA expression between FAC-treated and untreated cells. (**B**) Volcano plot showing overall changes in the mRNA expression between "Lip-1+FAC" and "MT+FAC"-treated cells.

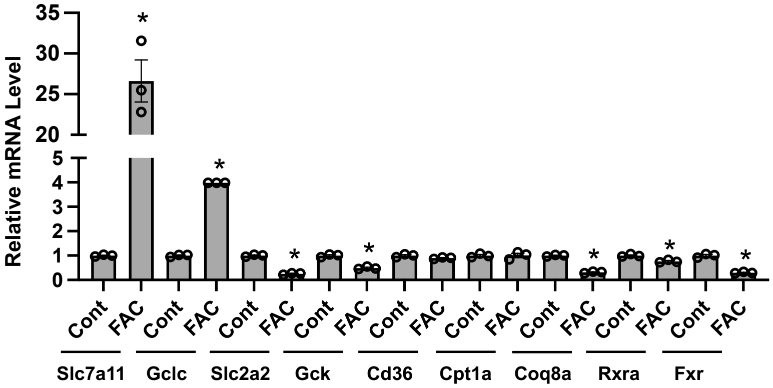

**Figure S2: The effects of iron overload on the expression of genes related to glutathione and CoQ biosynthesis, glycolysis, and fatty acid oxidation**. PMH were treated for 24 h with 100 μM FAC, followed by qRT-PCR analysis of transcript levels of the listed genes. Data shown are mean±SEM (n = 3). *, $P < 0.01$ compared to the untreated Cont cells.

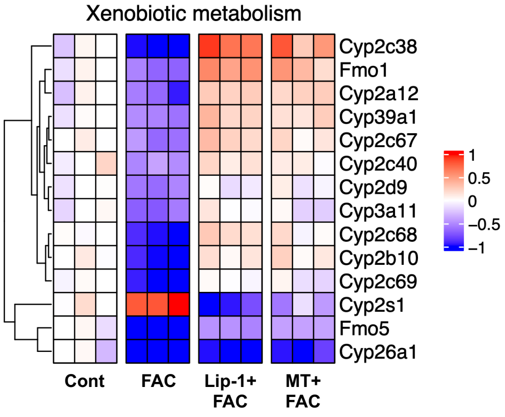

**Figure S3: The effects of iron overload and antioxidant treatment on the expression of genes in xenobiotic metabolism pathway**. The heatmap displays genes that were differentially expressed in PMH in response to iron overload and/or pretreatment with Lip-1 or MT.

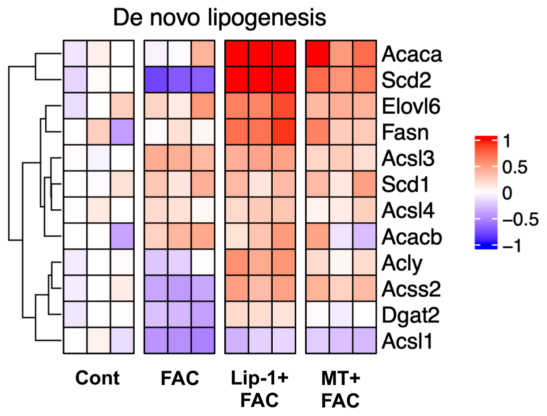

**Figure S4: The effects of iron overload and antioxidant treatment on the expression of genes that mediate *de novo* lipogenesis**. The heatmap displays genes that were differentially expressed in PMH in response to iron overload and/or pretreatment with Lip-1 or MT.

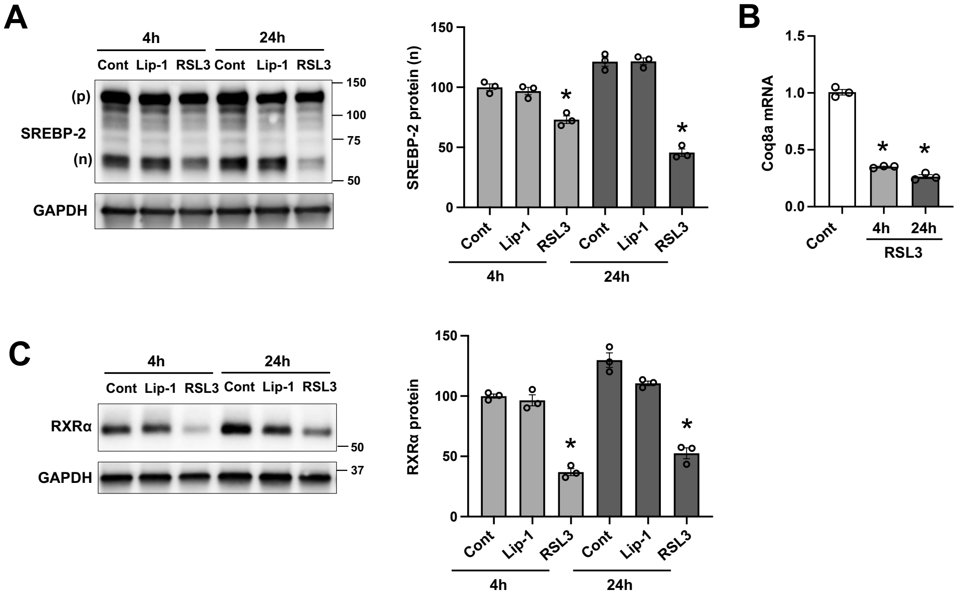

**Figure S5: Ferroptosis inducer RSL3 downregulates genes that mediate CoQ biosynthesis**. HepG2 cells were treated with 100 nM RSL3, a potent inducer of ferroptosis, or 5 μM Lip-1, a lipid ROS scavenger. (**A**) The expression of precursor (p) and nuclear (n) forms of SREBP-2 protein was determined by Western blotting. (**B**) Coq8a mRNA expression was determined by qRT-PCR. (**C**) RXRα protein level was determined by Western blotting. Data shown are mean±SEM (n = 3). *, *P* < 0.01 compared to the untreated Cont cells within the same group.

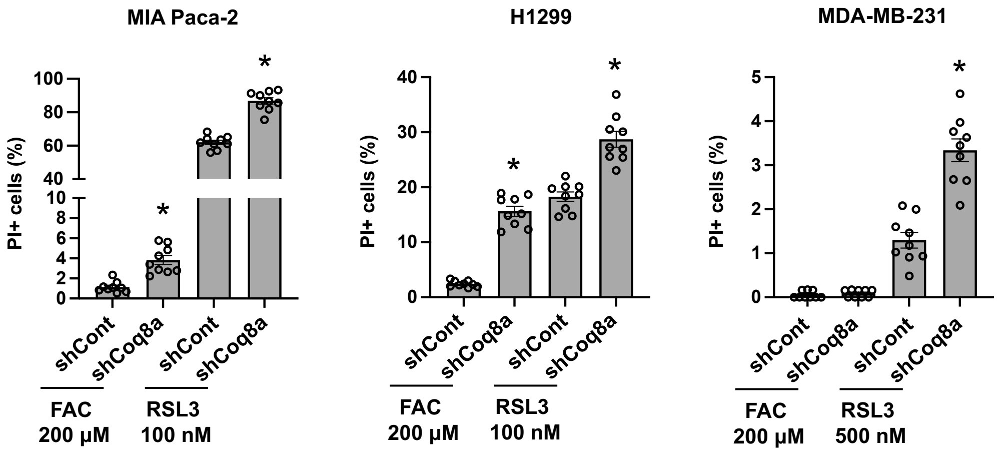

**Figure S6: CoQ8A deficiency promotes ferroptosis of various cancer cell lines**. Pancreatic cancer MIA Paca-2 cells, lung cancer H1299 cells, and breast cancer MDA-MB-231 cells, with or without the stable knockdown of CoQ8A, were treated for 4 h with the indicated concentrations of FAC or RSL3. Ferroptotic cells were stained with 2 μg/ml propidium iodide (PI), and the percentage of PI+ cells are shown as mean±SEM (n = 9 fields of 3 independent experiments). *, $P < 0.01$ compared to control cells within the same treatment group.

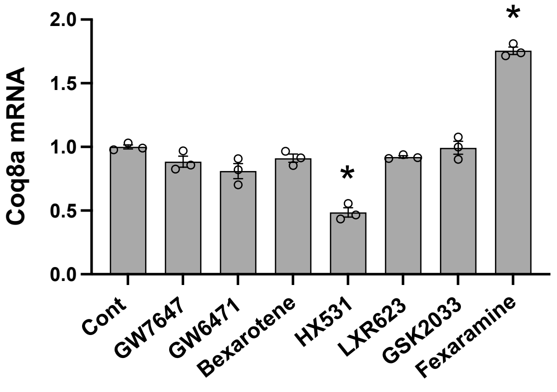

**Figure S7: The effects of nuclear receptor activation or inhibition on Coq8a expression**. HepG2 cells were treated for 24 h with pharmacological drugs GW7647 (5 μM), GW6471 (5 μM), bexarotene (2 μM), HX531 (5 μM), LXR623 (5 μM), GSK2033 (5 μM), or fexaramine (2 μM). Cells were then harvested for qRT-PCR analysis of Coq8a mRNA expression. Data shown are mean±SEM (n = 3). *, $P < 0.01$ compared to the untreated Cont cells.

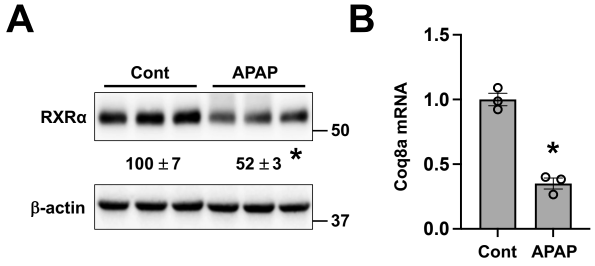

**Figure S8: Downregulation of RXRα and CoQ8A in acetaminophen induced liver injury**. Wild-type male mice were treated with 500 mg/kg acetaminophen (APAP). After 6 h, liver was harvested for the analysis of RXRα protein expression (**A**) and Coq8a mRNA expression (**B**) by Western blotting and qRT-PCR, respectively. *, $P < 0.01$ compared to the untreated control mice.

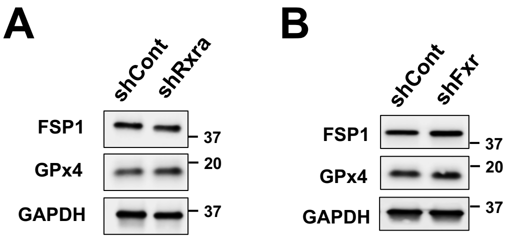

**Figure S9: GPx4 and FSP1 expression was not regulated by RXRα and FXR in hepatocytes**. The protein expression of GPx4 and FSP1 was determined in HepG2 cells with or without the knockdown of RXRα (**A**) or FXR (**B**). Shown are representative blots from three independent experiments.
